# *De novo* recovery of Ghana virus, an African bat Henipavirus, reveals differential tropism and attenuated pathogenicity compared to Nipah virus

**DOI:** 10.1101/2025.10.08.679836

**Authors:** Griffin D. Haas, Olivier Escaffre, Rebecca A. Reis, Terry L. Juelich, Jennifer K. Smith, Lihong Zhang, Birte K. Kalveram, Axel A. Guzmán-Solís, Dariia Vyshenska, William Klain, Alexander L. Greninger, Alexander N. Freiberg, Benhur Lee

## Abstract

Henipaviruses (HNVs) like Nipah (NiV) and Hendra (HeV) viruses represent severe zoonotic threats. Ghana virus (GhV), identified in 2012, is the only African bat henipavirus with a near-complete genome assembly. However, without isolates in culture, GhV biology, pathogenicity, and zoonotic potential remain poorly understood. Using reverse genetics, we recovered a full-length infectious clone of GhV at BSL-4 following rational reconstruction of its incomplete 3′ leader and modification of a non-canonical transcriptional initiation site. GhV demonstrated restricted receptor tropism (ephrin-B2 but not ephrin-B3) and distinct innate immune antagonism. Replication was attenuated in primary human cells, but was enhanced in bat cells. In Syrian golden hamsters, GhV infection caused no disease or mortality. Furthermore, a chimeric NiV encoding the GhV receptor-binding protein was completely attenuated *in vivo*, implicating ephrin-B3 receptor usage as a critical determinant of HNV pathogenesis. These findings elucidate GhV zoonotic potential and inform strategies for virus surveillance and control.

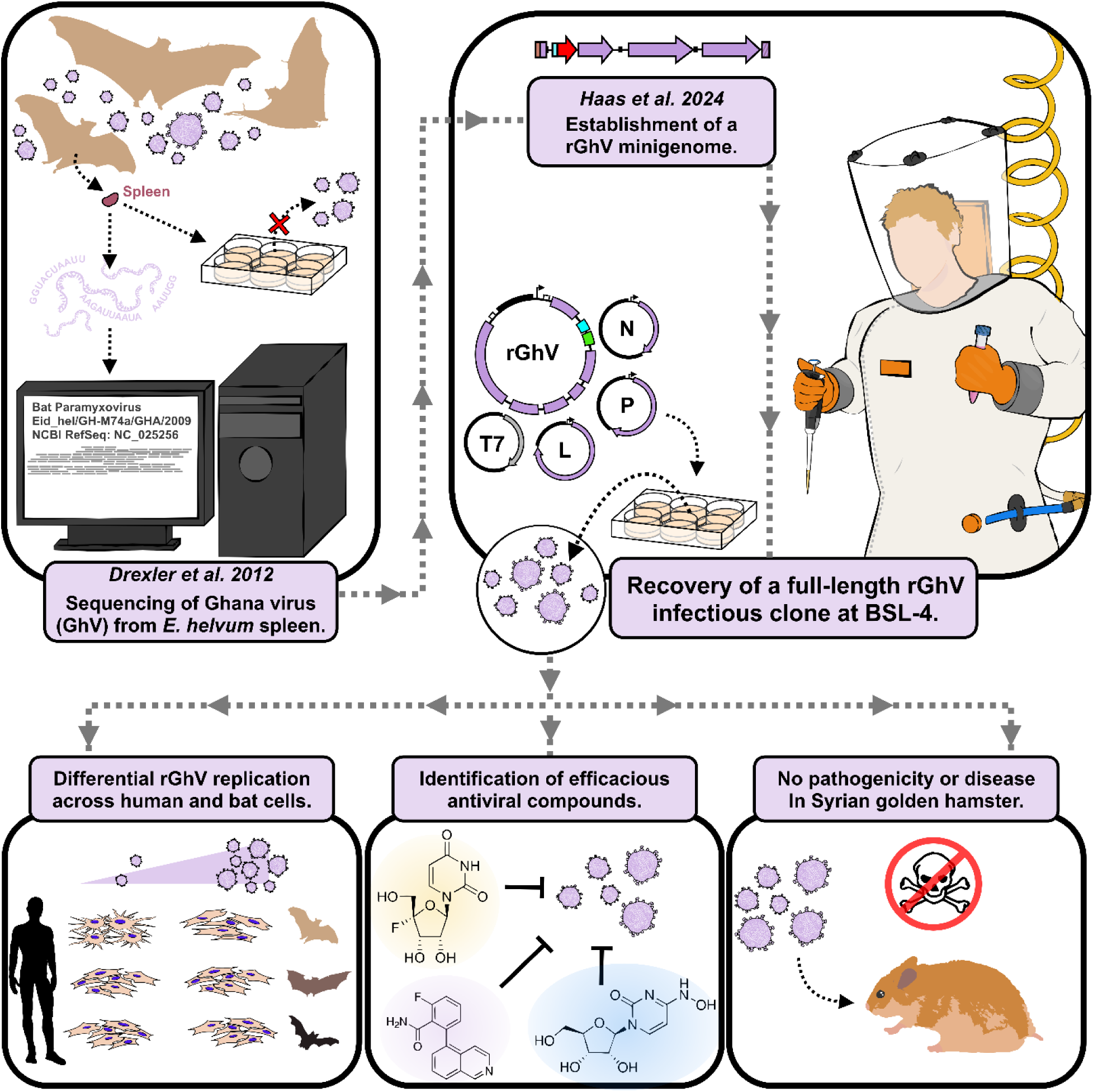

## Introduction

Bats are reservoirs of paramyxoviruses, including highly pathogenic henipaviruses (HNVs) such as Nipah virus (NiV) and Hendra virus (HeV)^1–4^. Spillover events of NiV and HeV have been geographically restricted to Southeast Asia and Australia, corresponding to the range of their reservoir host, Pteropid (*Pteropus* genus) bats^5,6^. However, recent wildlife surveillance efforts have made it evident that HNVs are globally distributed, with significant evidence of HNV circulation in African fruit bats. Metagenomic and serological surveys of *Eidolon helvum* (*E. helvum*) bats in Africa have consistently identified evidence of HNV infection^7–12^. Given that an estimated 128,000 *E. helvum* bats are captured annually as bushmeat in Ghana alone, these findings have raised concerns regarding the potential for African bat HNVs to spill over into humans^13^. In a cohort from Cameroon, seropositivity to HNVs was exclusively observed in individuals involved in butchering bats, suggesting that such spillover events are already occurring^11^.

In 2012, a novel HNV genome was identified from the spleen tissue of an *E. helvum* bat, leading to the discovery of Ghana virus (GhV, strain M74a)^1^. To date, GhV remains the only available genome of an African bat HNV. However, GhV is known by its sequence alone, and has yet to be isolated in culture. The dearth of isolates for GhV has hindered our understanding of its biology, pathogenicity, and potential risk to human health. We recently demonstrated that the GhV replicase is functional in chimeric minigenome assays, suggesting that reverse genetics approaches can be applied to GhV^14^.

A combination of receptor usage and potent antagonism of the host innate immune response is thought to underlie the severe disease caused by prototypical HNVs^15,16^. NiV and HeV use the highly-conserved ephrin-B2 (EFNB2) and ephrin-B3 (EFNB3) receptors to enter cells^17–19^. EFNB2 is more ubiquitously expressed across neurons, endothelial cells, and the smooth muscle of arterial vessels, while EFNB3 expression is largely restricted to the central nervous system; the tissue distribution of EFNB2 and EFNB3 corresponds to the tropism and neurovirulence of NiV and HeV^20,21^. By contrast, experimental binding and structural studies suggest that GhV only utilizes EFNB2^22^. However, the *in vivo* consequences of this more restricted receptor usage have not been explored due to the lack of viral isolates. Furthermore, NiV encodes multiple virulence factors that suppress host antiviral responses during infection^23,24^. It remains unknown whether the corresponding orthologs in GhV are expressed or are functionally sufficient to antagonize the human innate immune response.

In this study, we used reverse genetics to recover a full-length, infectious clone of GhV *de novo* at BSL-4. Recovery of GhV required modification of a non-canonical gene start motif upstream of the M gene. GhV replication was attenuated relative to NiV in primary human and porcine cells, but certain bat cell lines supported more robust GhV replication; furthermore, a subset of bat cells supported better replication of GhV than NiV. Experimental infection of Syrian golden hamsters with GhV resulted in no disease or mortality. Furthermore, we demonstrate that EFNB3 receptor usage is a major determinant of HNV pathogenesis by rescuing a recombinant NiV (rNiV) chimera encoding the GhV receptor binding protein, which was completely attenuated *in vivo*. This work provides the first experimental characterization of full-length, replication competent GhV in culture and establishes a foundation for further studies into its biology, host range, pathogenicity, and zoonotic potential.

## Results

### The gene start encoded upstream of the matrix gene in GhV is non-canonical and does not drive protein expression in a GhV minigenome system

We previously demonstrated that the available genome sequence of GhV M74a is incomplete, lacking 28 nucleotides at the extreme 3′ terminal end spanning its genomic promoter. However, substitution of this missing sequence with the homologous promoter element from HeV was sufficient to facilitate recognition by the GhV RNA-dependent RNA polymerase (vRdRp) in chimeric minigenome assays^14^. Building upon this approach, we constructed and rescued a full-length rGhV reverse genetics system encoding a *Gaussia* luciferase and eGFP reporter cassette between the N and P genes **(Figure S1A)**. Initial attempts to rescue this virus produced GFP positive foci by 11 days post-transfection (DPT); however, by 20 DPT, expansion of these foci was minimal, and the spread appeared to be predominantly driven by cell-to-cell contact, producing few if any distal infection events **(Figure S1B)**. This suggested that the reference sequence of GhV was unable to achieve efficient viral particle spread.

Having confirmed through comprehensive *in silico* analysis that the coding sequences (CDS) of each gene in GhV were consistent with the expected HNV gene architecture **(Figure S2)**, we next examined the reference genome for potential assembly variations in intergenic regions that might affect viral replication efficiency. To achieve expression, each viral gene must be flanked by an upstream gene start (GS) sequence and a downstream gene end (GE) sequence. Recognition of a GS sequence triggers the vRdRp to initiate transcription and capping of a nascent viral mRNA, whereas recognition of a GE sequence triggers the vRdRp to polyadenylate the nascent strand and terminate transcription^25–27^. Sequence alignment of bat-borne HNV GS sequences demonstrated an absolutely conserved 3′-UCCU-5′ motif localized immediately downstream of each intergenic region (IGR) for viral genes, while GE sequences consisted of a highly conserved 3′-AAUNNNUUUU-5′ motif. Based upon these consensus motifs, we were able to identify putative GS and GE sequences flanking each GhV gene (**Figure S3A**). Curiously, the GS upstream of the matrix gene (GhV-M) began with the motif 3′-UCC<u>C</u>-5′, and was the sole GS observed to break the otherwise absolutely conserved 3′-UCCU-5′ consensus **(Figure 1A, Figure S3A)**.

**Figure 1.**
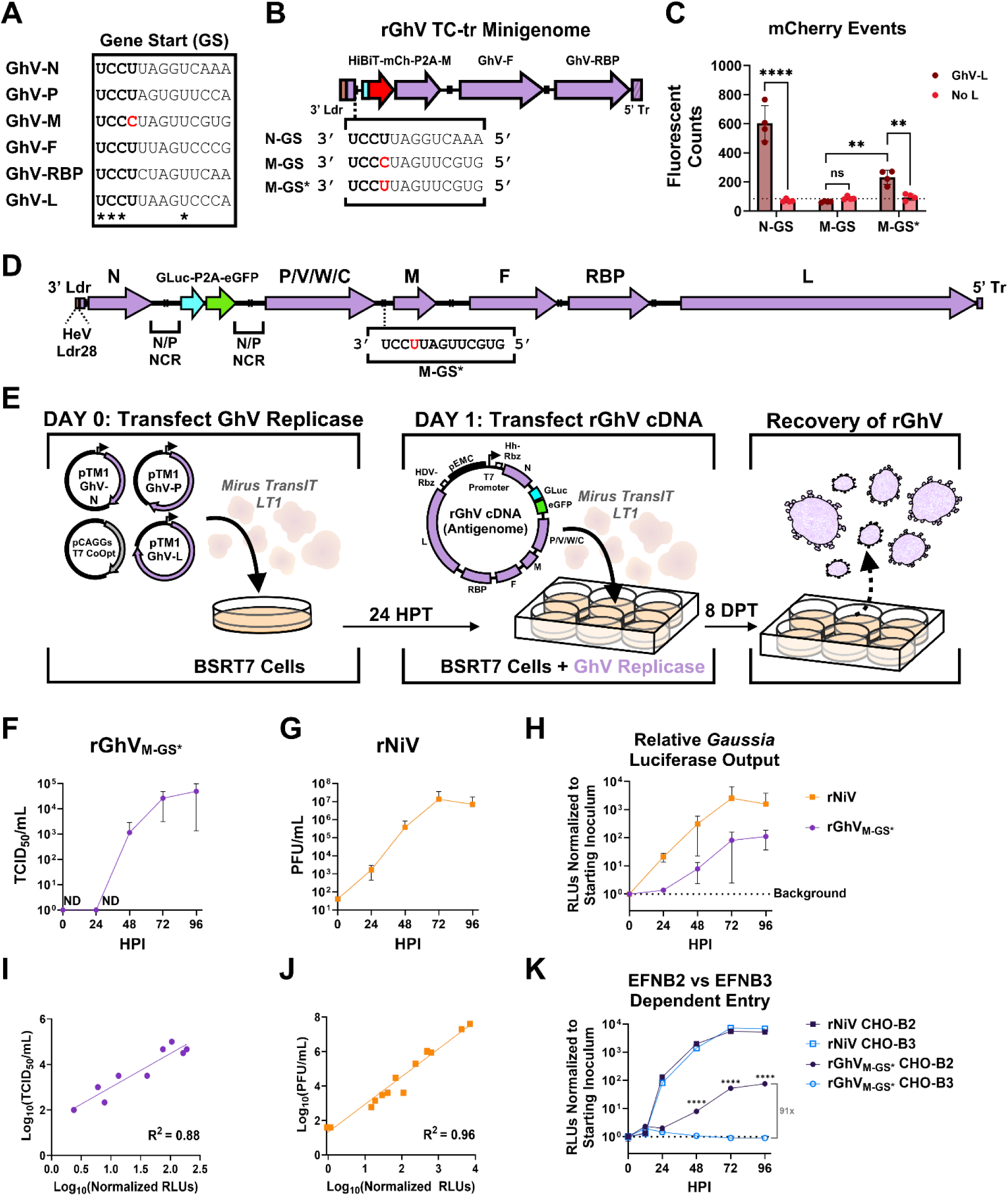
Restoring a canonical gene start in GhV strain M74a enables rescue and amplification of rGhV *de novo*. **(A)** Alignment of the gene start (GS) sequences encoded upstream of each gene in GhV M74a. The ‘UCCU’ consensus motif is shown in bold, and absolutely conserved bases are denoted with an asterisk below the alignment. The non-canonical base in the GhV-M GS is highlighted in red. **(B)** Design of rGhV TC-tr minigenomes in which the HiBiT-mCherry reporter gene is encoded under the control of either the N-GS, M-GS, or a modified M-GS (M-GS*), respectively; the GS sequences controlling HiBiT-mCherry expression are shown next to each label, and the non-canonical base that is changed in the M-GS to make M-GS* is highlighted in red. **(C)** Quantification of mCherry-positive events resulting from replication and transcription of each rGhV TC-tr minigenome either in the presence or absence of GhV-L. Matched microscopy and western blot data are presented in **Figure S3B** and **S3C**. **(D)** Schematic of the rGhV_M-GS*_ genome, in which the M-GS was modified to be consensus-like. A *Gaussia* luciferase eGFP reporter gene cassette is accommodated by duplication of the NCR between the N and P genes. The unmapped 28 nucleotides of the GhV M74a genomic promoter sequence were replaced with the homologous genomic promoter sequence of Hendra virus (HeV Ldr28). **(E)** Schematic detailing the rescue approach of rGhV, in which BSRT7 cells are transfected with GhV-N, -P, -L, and T7 polymerase on day 0, and are subsequently transfected with the antigenomic plasmid at day 1. Cells are then monitored for reporter gene signal and virus amplification. **(F)** Growth of rGhV_M-GS*_ in CHO-B2 cells at an MOI of 0.05. Supernatants were harvested with replacement at 0, 24, 48, 72, and 96 HPI and used for virus titration (TCID_50_/mL). ND indicates no detectable virus. **(G)** Growth of rNiV in CHO-B2 cells at an MOI of 0.05. were harvested with replacement at 0, 24, 48, 72, and 96 HPI and used for virus titration (PFU/mL). **(H)** Corresponding *Gaussia* luciferase activity from matched supernatants from (F) and (G), with RLUs normalized to the signal from the 0 HPI timepoint for each respective virus. Correlation between log_10_(titer) and log_10_(normalized RLUs) for rGhV_M-GS*_ (purple) **(I)** and rNiV (orange) **(J)** was used to determine the reliability of *Gaussia* luciferase as a surrogate for virus replication. **(K)** Relative replication of rNiV (squares) and rGhV_M-GS*_ (circles) in CHO-B2 cells (dark blue) and CHO-B3 cells (light blue). CHO-B2 and CHO-B3 cells were infected at an MOI of 0.05 and supernatants were harvested at 0, 12, 24, 48, 72, and 96 HPI for *Gaussia* luciferase assays. RLUs were normalized to the starting inoculum (0 HPI) for each respective virus. Statistical significance was determined by two-way ANOVA with Šídák’s multiple comparisons test in GraphPad Prism to compare relative infection in CHO-B2 and CHO-B3 at each timepoint. Brackets to the right of the graph in (K) denote fold-change between RLUs generated from infection of CHO-B2 and CHO-B3 cells. Non-significant datapoints are unlabeled (P > 0.05); *, P ≤ 0.05; **, P ≤ 0.01; ***, P ≤ 0.001; ****, P ≤ 0.0001. All experiments were conducted in biological triplicate and error bars represent standard deviation.

Because the matrix protein mediates viral particle production, we sought to investigate if its non-canonical motif impacted GS activity in our previously described rGhV tetracistronic, transcription and replication competent (TC-tr) minigenome^14^. We generated three separate constructs in which the HiBiT-mCherry reporter gene was encoded under the control of the GhV-N GS, the reported native GhV-M GS, or a modified GhV-M GS (denoted as M-GS*) in which we made a single nucleotide change to restore the 3′-UCCU-5′ consensus motif (**Figure 1B**). These respective rGhV TC-tr minigenomes were co-transfected into BSRT7 cells along with plasmids encoding codon-optimized T7 polymerase, T7-driven GhV-N and GhV-P, and either GhV-L or an irrelevant protein (GFP) as a no-L control. The GhV-N GS was sufficient to drive significant expression of the HiBiT-mCherry reporter gene by the vRdRp, yielding several hundred mCherry events over the no-L control (**Figure 1C**). However, when the M-GS was employed, we observed no mCherry positive cells over the no-L control. Notably, the single-nucleotide change implemented in the minigenome encoding M-GS* was sufficient to restore partial activity, resulting in significant, albeit modest expression of the HiBiT-mCherry reporter gene over the no-L control **(Figure 1C, S3B and S3C)**.

### A rGhV encoding a M-GS bearing the consensus motif (M-GS*) can be rescued and amplified in culture, albeit with delayed kinetics relative to NiV

Based on our observation that the native GhV M-GS does not drive protein expression in the minigenome context, we hypothesized that the initial rGhV rescue failed to amplify in culture due to a deficiency in GhV-M expression. We encoded a single nucleotide change in the M-GS in our full-length rGhV reverse genetics plasmid, such that the GhV-M gene was instead under control of the functional M-GS* **(Figure 1D)**. BSRT7 cells were co-transfected with plasmid encoding codon optimized T7 polymerase and plasmids encoding the GhV replicase (-N, -P, and -L genes) under T7 promoter. Twenty-four hours later, these cells were transfected again at high containment with plasmid encoding the rGhV antigenome with M-GS* (rGhV_M-GS*_) (**Figure 1E)**. By 8 DPT, GFP positive foci were observed throughout the rescue well, with obvious instances of particle spread and cell-to-cell spread **(Figure S3D)**. Following amplification, stocks of rGhV_M-GS*_ were validated by RNA sequencing and titrated by TCID50 on highly permissive Chinese hamster ovary cells that over-express EFNB2 (CHO-B2), the HNV entry receptor. Notably, RNA sequencing confirmed that the HeV genomic promoter, used in lieu of the unmapped GhV genomic promoter, remained unchanged during virus amplification.

To assess replication kinetics, CHO-B2 cells were infected at an MOI of 0.05 with either rGhV_M-GS*_ or rNiV (Malaysia strain), with both viruses encoding a GLuc-P2A-eGFP reporter cassette. Cell supernatants were collected at 0, 24, 48, 72, and 96 hours post-infection (HPI) for virus titration and *Gaussia* luciferase assay. rGhV_M-GS*_ showed exponential growth and achieved an endpoint titer of 5×10^4^ TCID₅₀/mL by 96 HPI with an eclipse phase of >24 HPI **(Figure 1F)**. By contrast, rNiV infection of CHO-B2 cells yielded measurable progeny virus as early as 24 HPI **(Figure 1G)**. The delayed kinetics for rGhV_M-GS*_ mirror observations made using TC-tr minigenomes, in which the GhV replicase failed to generate appreciable reporter gene signal until at least 48 HPT, whereas the NiV replicase produced signal as early as 24 HPT **(Figure S4A and S4B)**. *Gaussia* luciferase production reflected the titration data at each timepoint **(Fig. 1H)**, demonstrating a strong positive correlation between RLUs and virus titer for both rGhV_M-GS*_ **(Fig 1I)** and rNiV **(Fig 1J)**. Thus, *Gaussia* luciferase serves as a reliable surrogate for measuring viral replication.

We previously demonstrated that GhV utilizes EFNB2, but not EFNB3, for entry into cells^22^. To validate GhV receptor usage in full-length virus, CHO cells expressing either EFNB2 (CHO-B2) or EFNB3 (CHO-B3) were infected with either rGhV_M-GS*_ or rNiV at an MOI of 0.05, and supernatant was collected at 0, 12, 24, 48, 72, and 96 HPI for *Gaussia* luciferase assay. While rNiV could replicate equally well in both CHO-B2 and CHO-B3 cells, rGhV_M-GS*_ could only replicate in CHO-B2 cells (91-fold increase) with no obvious infection or amplification in CHO-B3 cells **(Figure 1K)**. Infection with rNiV resulted in widespread cytopathic effect (CPE) and the formation of large syncytia, whereas rGhV_M-GS*_ infection yielded predominantly single-cell infection events, with only occasional, modest syncytia **(Figure S5A-D)**.

### Co-transcriptional editing of the GhV-P gene yields products that antagonize human innate immune signaling in a manner distinct from NiV

To characterize the transcriptional profile of GhV, we employed nanopore long-read direct RNA sequencing on total RNA from rGhV_M-GS*_ infected CHO-B2 cells at 72 HPI. Mapping of mRNA reads to the rGhV_M-GS*_ genome revealed that during infection, the vRdRp produces a characteristic 3′-5′ transcriptional gradient, where N > P > M > F > RBP > L **(Figure S6A)**. Furthermore, this allowed us to map the GS and GE sequences of GhV, confirming our predicted gene boundaries in **Figure 1** and **Figure S3A**.

During infection, the prototypical HNVs generate potent antagonists of innate immune signaling by co-transcriptional editing of the phosphoprotein gene^23,24^. These antagonists, ‘V’ and ‘W’, are generated when the vRdRp inserts one or two non-templated guanosine nucleosides, respectively, into nascent P-gene transcripts at a characteristic editing site. The GhV-P gene encodes such an editing site and has the coding repertoire to make V and W proteins (**Figure 2A**). Analysis of reads mapping to the editing site revealed that co-transcriptional editing does occur during GhV infection, with 79% of transcripts encoding P, 13% encoding V, and 8% encoding W **(Figure S6B and S6C)**. This proportion of P-edited transcripts (V+W=21%) appears lower than the >50% of P-edited transcripts found in NiV and HeV^23,28,29^.

**Figure 2.**
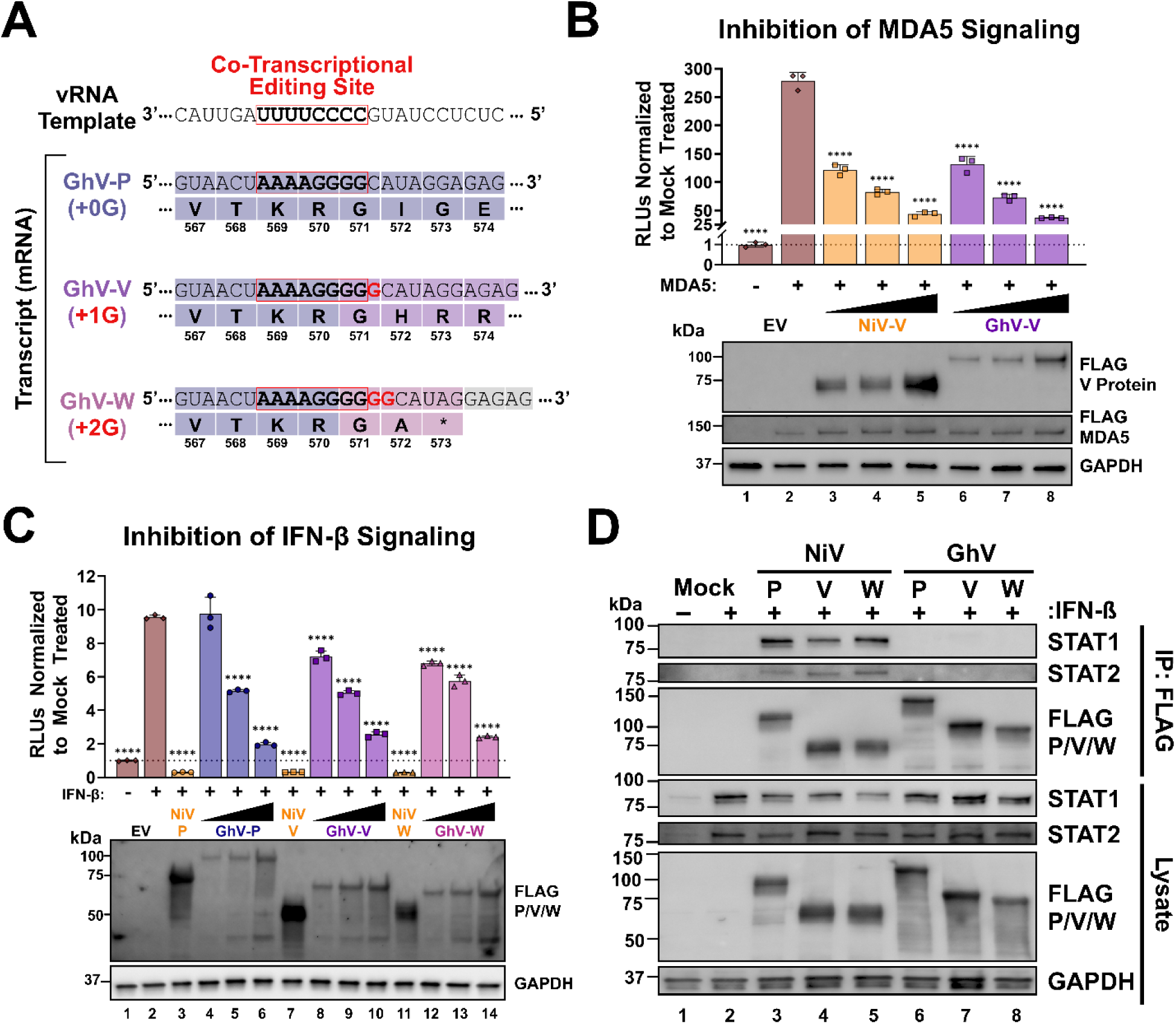
Co-transcriptional editing of the GhV-P gene yields products that antagonize human innate immune signaling in a manner distinct from NiV. **(A)** Schematic of the co-transcriptional editing site encoded in the GhV-P gene. Insertion of a single non-templated guanosine nucleoside at the editing site yields GhV-V, and the insertion of two non-templated guanosines yields GhV-W. The mRNA sequence for each gene product is shown above its respective coding sequence, with non-templated nucleosides depicted in red. **(B)** Inhibition of MDA5-driven signaling by NiV-V and GhV-V in HEK-293T cells. A FLAG-tagged MDA5 expression plasmid, pISRE-TA-Luc, and pRL-TK were co-transfected with either EV or increasing amounts (50ng, 100ng, or 200ng) of NiV-V or GhV-V. Dual-luciferase assays were performed at 24 HPT, and RLUs were normalized to the EV condition without MDA5 induction. **(C)** Inhibition of IFN-β– driven signaling by NiV-P, -V, and -W and GhV-P, -V, and -W. HEK-293T cells were co-transfected with pISRE-TA-Luc and pRL-TK with either EV or increasing amounts (50ng, 100ng, or 200ng) of respective viral protein. At 24 HPT, cells were treated with 1000 U/mL of human IFN-β for 6 hours. Dual-luciferase assays were then performed, and RLUs were normalized to the untreated EV control. **(D)** Co-immunoprecipitation of endogenous STAT1 and STAT2 by either FLAG-tagged NiV-P, -V, and -W or GhV-P, -V, and -W. Transfected HEK-293T cells treated with 1000 U/mL IFN-β, lysed, and subjected to immunoprecipitation (IP) using anti-FLAG beads. Western blots of the IP eluates and whole-cell lysate were then probed to detect viral proteins, STAT1, and STAT2. All experiments were conducted in biological triplicate, with error bars representing standard deviation. The Western blots in (B) and (C) confirm expression of the indicated proteins in each experimental condition, with GAPDH serving as a loading control. Statistical significance in (B) and (C) was assessed by one-way ANOVA with Dunnett’s multiple comparisons test, comparing each condition to the respective EV control. Non-significant data points (P > 0.05) are not labeled; *, P ≤ 0.05; **, P ≤ 0.01; ***, P ≤ 0.001; ****, P ≤ 0.0001.

To determine if the P-editing products of GhV are capable of antagonizing human innate immune signaling, we cloned FLAG-tagged P, V, and W proteins from NiV and GhV into a pCAGGS expression vector. Dual luciferase innate immune response reporter assays demonstrated that both GhV-V and NiV-V potently inhibited IFN-β promoter activation of innate immune signaling via MDA5, corroborating *in silico* predictions of MDA5 binding **(Figure 2B and Figure S6D)**. We further observed that the GhV-P, -V, and -W proteins drove a dose-dependent inhibition of transcription from an interferon stimulated response element (ISRE) promoter in the presence of IFN-β (**Figure 2C**). To determine if the mechanism of IFN-β signaling antagonism is conserved between NiV and GhV, we pulled down FLAG-tagged P, V, and W proteins and blotted for human STAT1 and STAT2. While we detected co-immunoprecipitation of human STAT1 and STAT2 by NiV-P, -V, and -W, there was no detectable pull down of these factors by GhV-P, -V, and -W **(Figure 2D)**. The described STAT1-binding motif in NiV-P is poorly conserved in GhV-P, supporting the conclusion that GhV antagonism occurs in a manner independent of STAT1 binding **(Figure S6E)**.

### Primary human and porcine cells support limited replication of rGhV_M-GS*_

To better understand the zoonotic potential of GhV, we assessed its ability to replicate in a panel of primary human and porcine cells that were previously established to support robust rNiV replication. rNiV replicated more robustly than rGhV_M-GS*_ in primary human astrocytes (HAs) achieving >2000-fold increase in normalized RLUs (nRLU) compared to less than 10-fold increase in nRLUs for rGhV_M-GS*_ by 96 HPI **(Figure 3A)**. Nonetheless, rGhV_M-GS*_ clearly infected HAs and numerous small syncytia were evident (**Figure S7A**). In contrast, while rNiV replicated efficiently in both primary human brain microvascular endothelial cells (HBMECs, 775-fold increase in RLUs) and human umbilical vein endothelial cells (HUVECs, 499-fold increase in RLUs), rGhV_M-GS*_ showed no detectable signs of infection, and was apparently restricted in these cells **(Figure 3B and 3C)**. Similarly, albeit to a lesser extent, normal human bronchial epithelial cells (NHBEC, **Figure 3D)** and small airway epithelial cells (SAEC, **Figure 3E)** supported robust rNiV replication (81-fold and 78-fold increases in RLUs, respectively), but limited rGhV_M-GS*_ replication (∼2-fold increase in RLUs). Primary porcine kidney epithelial cells (PPKECs), however, supported both rNiV replication (1066-fold increase in RLUs) and moderate rGhV_M-GS*_ replication (17-fold increase in RLUs) **(Figure 3F)**. Fluorescence microscopy data for EGFP+ cells corroborating these results are presented in **Figure S7**.

**Figure 3.**
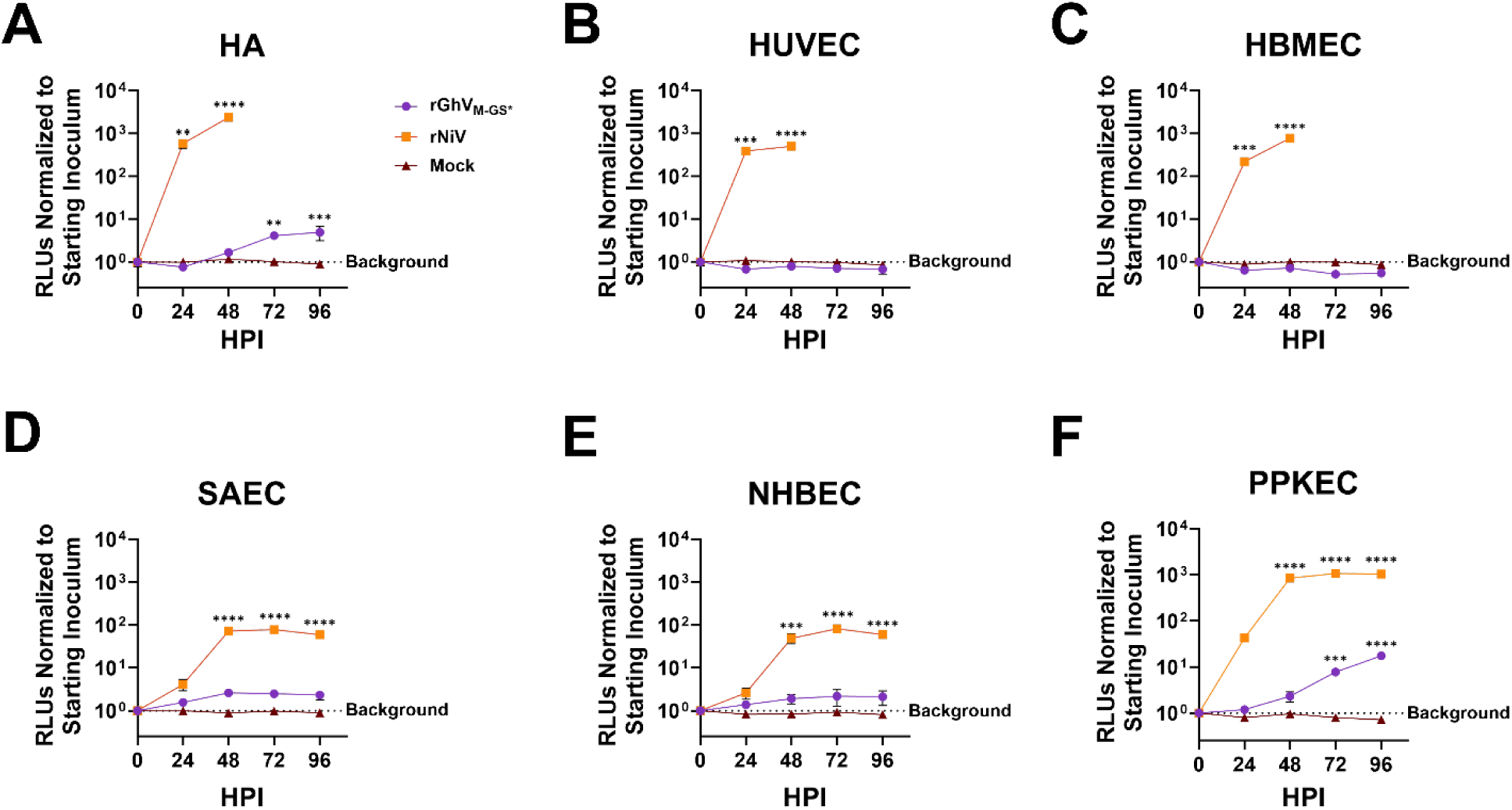
Primary human and porcine cells support limited rGhV_M-GS*_ replication. Replication kinetics of rGhV_M-GS*_ (purple), rNiV (orange), and mock-infected (brown) cell lines: **(A)** primary human astrocytes (HA); **(B)** human umbilical vein endothelial cells (HUVEC); **(C)** human brain microvascular endothelial cells (HBMEC); **(D)** small airway epithelial cells (SAEC); **(E)** normal human bronchial epithelial cells (NHBEC); and **(F)** primary porcine kidney epithelial cells (PPKEC). Cells were infected at an MOI of 0.1 and supernatants were collected with replacement at 0, 24, 48, 72, and 96 HPI. *Gaussia* luciferase activity (RLUs) in the supernatants were measured and normalized to the 0 HPI timepoint for each condition to measure increase in signal over time. Virus infections were conducted in at least biological triplicate and mock infections were conducted in at least biological duplicate. Statistical significance was determined by one-way ANOVA in GraphPad Prism with Dunnett’s multiple comparisons test to compare normalized RLUs at each time point to baseline RLUs at 0 HPI for each virus. Non-significant data points (P > 0.05) are not labeled; *, P ≤ 0.05; **, P ≤ 0.01; ***, P ≤ 0.001; ****, P ≤ 0.0001.

### A panel of bat cell lines supports differential rGhV and rNiV infection

Given the limited replication of rGhV_M-GS*_ across non-chiropteran cells, we next evaluated replication across a panel of bat-derived cell lines. These cells, representing five distinct bat species, were infected with either rNiV or rGhV_M-GS*_ using the same MOI (0.05) and monitored for signs of infection over the same amount time (96 HPI) as in **Figure 3**. The bat species included in this panel, their phylogeny, and geographical distribution are depicted in **Figure 4A and 4B**.

**Figure 4.**
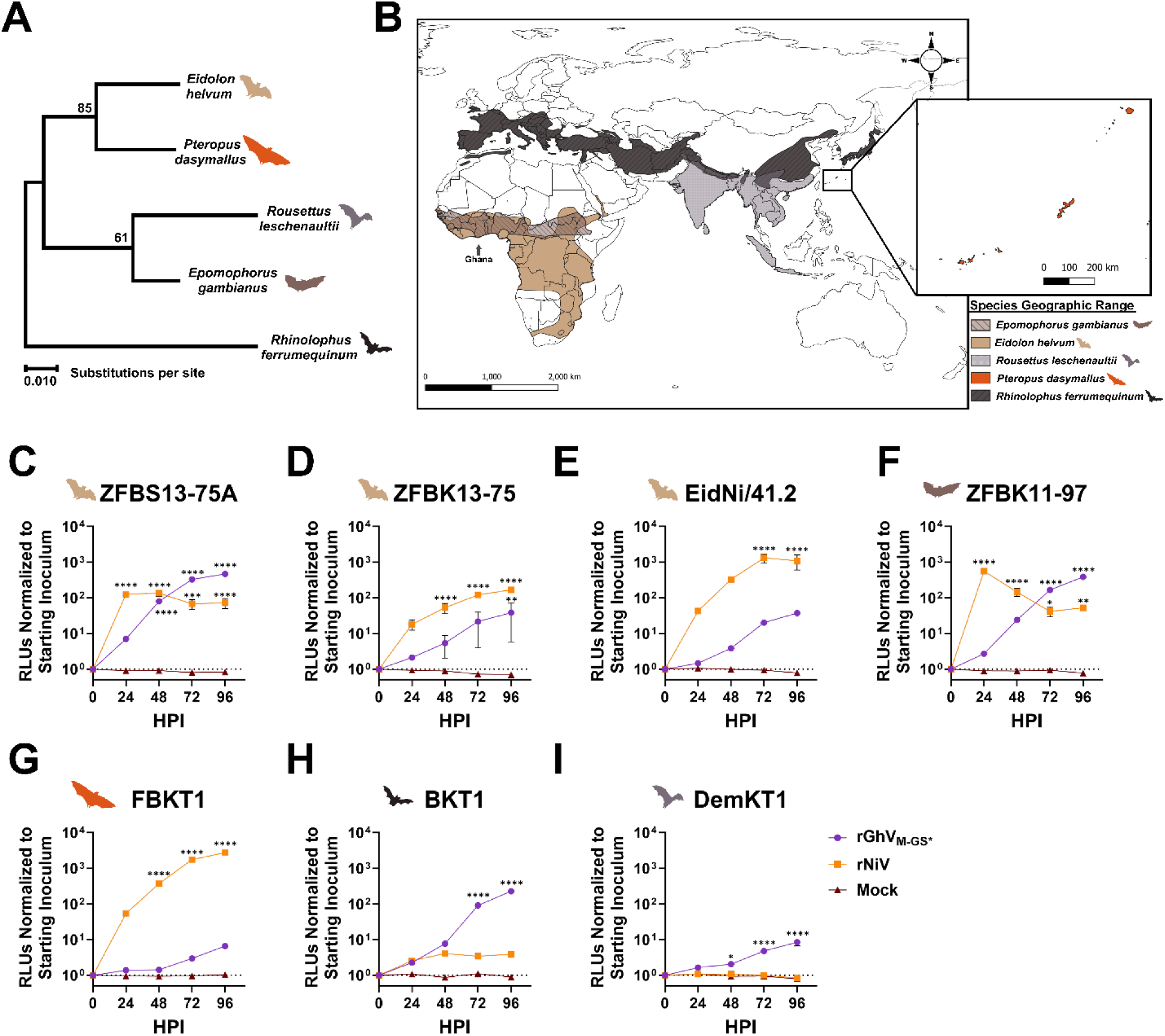
A panel of bat cells support differential replication of rGhV_M-GS*_ and rNiV. **(A)** Phylogenetic tree based on the cytochrome b gene from representative bat species in this study. Evolutionary history was inferred by using the Maximum Likelihood method and JTT matrix-based model in MEGA X, with the percentage of trees in which the associated taxa clustered indicated at the nodes. The tree is drawn to scale, with branch lengths representing amino acid substitutions per site. **(B)** Geographical range of respective bat species. Colored/shaded areas indicate species distribution ranges based on IUCN data, with *Eidolon helvum* depicted in tan; *Epomophorus gambianus* depicted in striped brown, *Pteropus dasymallus* depicted in orange (its range is shown in the inset); *Rousettus leschenaultii* depicted in light gray; and *Rhinolophus ferrumequinum* depicted with striped dark gray. The location of Ghana is marked with an arrow for geographic reference. Bat geographic ranges are courtesy of the IUCN Red List of Threatened Species (https://www.iucnredlist.org). Map projection was made using QGIS 3.40.7. Replication kinetics of rGhV_M-GS*_ (purple), rNiV (orange), or mock-infected (brown) bat cells were measured in: **(C) (**ZFB<u>S13-75A</u> cells, **(D)** ZFB<u>K13-75</u> cells, and **(E)** EidNi/41.2 cells from *Eidolon helvum* (straw-colored fruit bats); **(F)** ZFBK11-97 cells from *Epomophorus gambianus* (Gambian epauleted fruit bat); **(G)** FBKT1 cells from *Pteropus dasymallus* (Ryukyu flying fox); **(H)** BKT1 cells from *Rhinolophus ferrumequinum* (greater horseshoe bat); and **(H)** DemKT1 cells from *Rousettus leschenaultii* (Leschenault’s rousette). Cells were infected at an MOI of 0.05, and supernatants were harvested at 0, 24, 48, 72, and 96 HPI for GLuc assays. RLUs were normalized to the 0 HPI timepoint for each virus. Statistical significance was determined by two-way ANOVA in GraphPad Prism with Šídák’s multiple comparisons test to compare normalized RLUs at each time point with the corresponding RLUs at 0 HPI for each virus. Non-significant datapoints are unlabeled (P > 0.05); *, P ≤ 0.05; **, P ≤ 0.01; ***, P ≤ 0.001; ****, P ≤ 0.0001. All experiments were conducted in biological triplicate and error bars represent standard deviation.

*E. helvum* (straw colored fruit bat) is distributed broadly across sub-Saharan Africa and southward through Central Africa to Southern Africa **(Figure 4B)**. All *E. helvum* cells supported appreciable rNiV replication, demonstrating peak increases in nRLUs of 126-fold, 167-fold, and 1303-fold in ZFB<u>S13-75A</u> **(Figure 4C)**, ZFB<u>K13-75</u> **(Figure 4D)**, and EidNi/41.2 **(Figure 4E)**, respectively. In contrast to human and porcine cells, ZFB<u>S13-75A</u> cells also supported robust rGhV_M-GS*_ replication, yielding a 468-fold peak increase in nRLUs that was almost 4X higher than for rNiV **(Figure 4C)**. rGhV_M-GS*_ replication was more moderate in ZFB<u>K13-75</u> cells and in EidNi/41.2 cells, yielding 38-fold and 37-fold increases in nRLUs, respectively **(Figure 4D and 4E)**.

*Epomophorus gambianus* (Gambian epauletted fruit bat) is sympatric with *E.helvum*, as both bats inhabit regions across West and Central Africa, stretching from the Atlantic coast eastward into Central Africa **(Figure 4B)**. *E. gambianus* ZFBK11-97 cells supported robust replication by both rNiV and rGhV_M-GS*_, with 558-fold and 386-fold peak increases in nRLUs, respectively (**Figure 4F**). *Pteropus dasymallus* (Yaeyama fruit bat) is found mostly in East Asia and pteropodid bats are the reservoir species for NiV and HeV^5,6,30^ **(Figure 4B)**. *P. dasymallus* FBKT1 cells supported robust rNiV replication (2746-fold increase) with only minimal rGhV_M-GS*_ replication (7-fold) **(Figure 4G)**. Interestingly, two bat cell lines supported higher replication of rGhV_M-GS*_ than rNiV. *Rhinolophus ferrumequinum* (greater horseshoe bat) BKT1 cells supported more robust rGhV_M-GS*_ replication than NiV with a peak 227-fold versus 4-fold increase in nRLUs, respectively **(Figure 4H)**. The greater horseshoe bat, better known as the reservoir species for many coronaviruses^31–33^, has a wide geographical range that does not overlap with *E. helvum* **(Figure 4B)**. Finally, while *Rousettus leschenaultii* DemKT1 cells supported low-level replication of rGhV_M-GS*_ (8-fold increase in RLUs), rNiV replication was entirely restricted **(Figure 4I)**. While *R. leschenaultii* is found only in India and South-east Asia (**Figure 4B)**, *Rousettus spp.* are widespread and appear to host many RNA viruses with zoonotic potential (see Discussion)^34^.

To determine if receptor expression could partially account for the differential ability of these bat cell lines to support NiV versus GhV replication, we quantified the relative abundance of *EFNB2* transcripts from each bat cell line by RT-qPCR and plotted their relative expression levels against the peak nLuc values for GhV and NiV, respectively **(Figure S8A-B)**. While there appears to be a moderate correlation between relative *EFNB2* expression and peak nRLUs for GhV, this was not true for NiV **(Figure S8A-B).** *EFNB3* expression was uniformly low across these bat cells except for DemKT1 cells and is unlikely to play a role in the differential susceptibility patterns seen, especially since GhV does not use EFNB3 for entry^22^ **(Figure 1K)**. Fluorescence microscopy data largely corroborate the infection data in **Figure 4** (**Figure S8C-I)**.

### Experimental infection of Syrian golden hamsters with rGhV_M-GS*_ does not cause disease nor mortality

The Syrian golden hamster is a robust animal model for HNV disease and pathogenesis^35–37^. To evaluate the pathogenicity of GhV, we experimentally infected Syrian golden hamsters (n=6 per group, equal sex) with either 2.5 x 10^4^ TCID_50_ of rGhV_M-GS*_ or 2.5 x 10^4^ PFU of rNiV (back-titrated at 1.38 x 10^4^ PFU). Both viruses were administered either intraperitoneally (IP) or intranasally (IN). A mock-infected control group (n=4) was maintained in parallel. Animals were observed daily for clinical signs of illness and mortality over 21 days, and retro-orbital (RO) bleeds were collected at 2, 4, 6, and 8 DPI **(Figure 5A)**. As a surrogate for measuring virus replication *in vivo*, *Gaussia* luciferase activity was determined in serum from RO and terminal bleeds^11,38,39^. While hamsters infected with rNiV demonstrated a marked increase in RLUs relative to the mock control as early as 2 DPI, hamsters infected with rGhV_M-GS*_ exhibited no appreciable rise in RLUs at any timepoint **(Figure 5B)**. Consistent with this observation, hamsters infected with rGhV_M-GS*_ experienced no significant weight loss **(Figure 5C)** nor mortality **(Figure 5D)**. By contrast, animals infected with rNiV experienced significant weight loss between 4 and 8 DPI **(Figure 5C)**, with 6/6 animals succumbing to infection in the IN challenge group, and 5/6 succumbing to infection in the IP challenge group **(Figure 5D)**. To confirm that hamsters had been successfully infected with rGhV_M-GS*_, terminal sera collected at 21 DPI from survivors was tested in neutralization assays against rGhV_M-GS*_. Although animals challenged intranasally failed to seroconvert, 3/6 animals that received rGhV_M-GS*_ intraperitoneally developed neutralizing titers above the limit of detection (**Figure 5E**).

**Figure 5.**
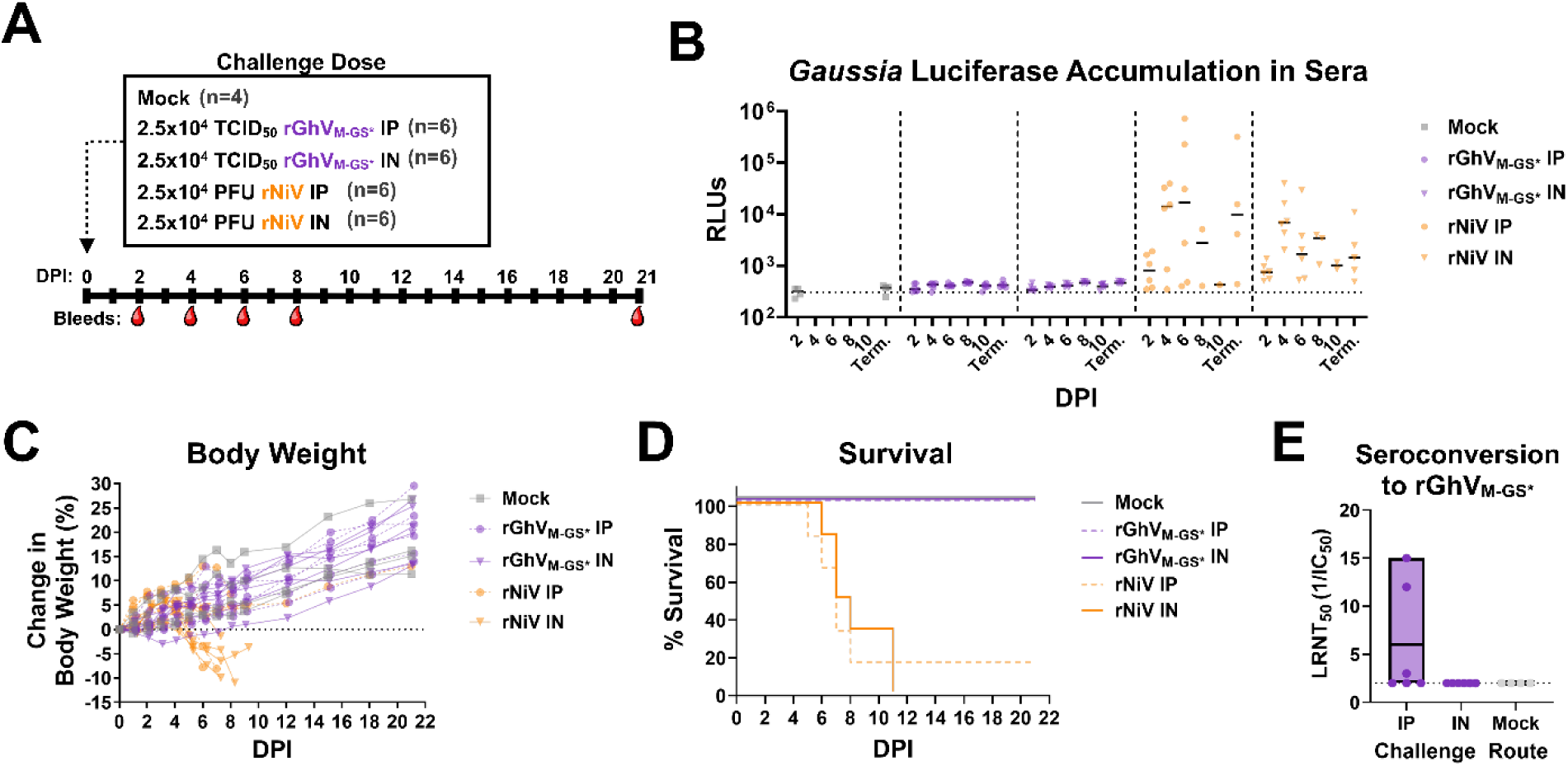
rGhV_M-GS*_ challenge in Syrian Golden Hamster animal model. **(A)** Schematic of the challenge design. Groups of Syrian golden hamsters were inoculated either intraperitoneally (IP) or intranasally (IN) route with 2.5×10^4^ TCID_50_ of rGhV_M-GS*_ (n=6 per group) or 2.5×10^4^ PFU of rNiV (n=6 per group). Mock infected (n=4) animals served as controls. Retro-orbital (RO) bleeds were collected at 2, 4, 6, and 8 DPI, with terminal bleeds collected at time of euthanasia or at 21 DPI. **(B)** Accumulation of *Gaussia* luciferase in serum as a surrogate for measuring virus replication in animals. Sera from RO and terminal bleeds was used for GLuc assays. **(C)** Percent change in body weight over time. **(D)** Survival curves of hamsters challenged with rGhV_M-GS*_ or rNiV compared to mock. **(E)** Quantification of luciferase neutralizing antibody titers against rGhV_M-GS*_ (LRNT_50_) in terminal sera from rGhV_M-GS*_ or mock-challenged animals. For all graphs, each datapoint represents an individual animal. Mock is depicted in gray (rectangles), rGhV_M-GS*_ is depicted in purple, and rNiV is depicted in orange. IP groups are depicted as circles, and IN groups are depicted as triangles.

### A chimeric, EFNB3-blind rNiV is completely attenuated in Syrian golden hamsters

To evaluate the role of differential receptor usage (EFNB2 vs EFNB3) in HNV pathogenicity, we constructed a chimeric NiV clone in which the NiV receptor binding protein (RBP) was replaced with the CDS of the GhV-RBP **(Figure S9A)**. We anticipated that this virus would be viable, as previous studies indicated that the GhV-RBP was capable of complementing NiV-F in fusion assay^40,41^. The chimeric virus, rNiV_GhV-RBP_, was rescued and amplified in culture, and demonstrated kinetics comparable to the isogenic WT rNiV **(Figure S9B)**. Furthermore, like GhV, the chimeric virus was unable to infect CHO-B3 cells **(Figure S9C)**. To determine the pathogenicity of the chimeric virus, Syrian golden hamsters (n=4 per group) were challenged intraperitoneally with either 1×10^4^ or 1×10^5^ PFU of rNiV_GhV-RBP_ (back-titrated at 2.23 x 10^4^ and 1.8 x10^5^ PFU) or WT rNiV (back-titrated at 1.1 x 10^4^ and 8.67 x 10^4^ PFU).

Animals were monitored for health and weight loss up until 21 DPI, with RO bleeds collected at 2, 4, 6, and 8 DPI. At 21 DPI, all survivors were bled, and at 28 DPI, surviving animals were re-challenged with 1×10^6^ PFU of WT rNiV to assess whether prior infection conferred protective immunity **(Figure 6A)**.

**Figure 6.**
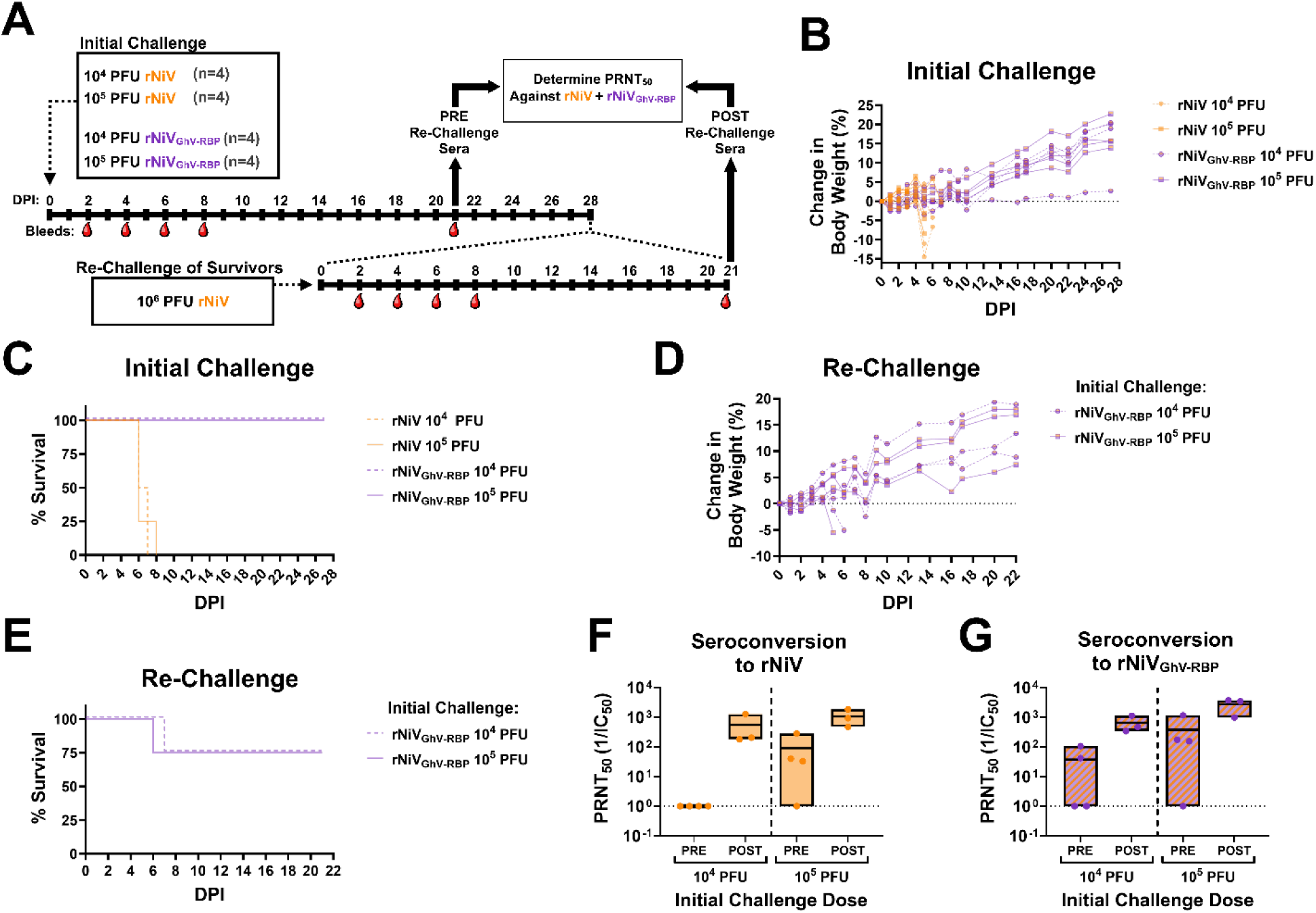
Syrian golden hamster challenge with rNiV and rNiV_GhV-RBP_. **(A)** Schematic of the two-phase animal challenge. Syrian golden hamsters (n=4 per group) were initially challenged with either 10^4^ PFU or 10^5^ PFU of rNiV or chimeric rNiV encoding the GhV-RBP (rNiV_GhV-RBP_). RO bleeds were collected at 2, 4, 6, 8, and 21 DPI. At 28 DPI, survivors were re-challenged with 10^6^ PFU of rNiV, with RO bleeds collected at 2, 4, 6, 8, and 21 days post re-challenge. Sera collected at 21 days post initial challenge (PRE re-challenge) and at 21 days post re-challenge (POST re-challenge) were used to determine seroconversion by PRNT_50_. **(B)** Percent change in body weight over time following the initial challenge with rNiV (orange) or rNiV_GhV-RBP_ (half-orange, half-purple). Each data point represents an individual animal; circles indicate a dose of 10^4^ PFU, and squares depict a dose of 10^5^ PFU. **(C)** Survival curves from the initial challenge. **(D)** Percent change in body weight over time following the re-challenge of survivors with 10^6^ PFU of rNiV. Datapoint depictions reflect the initial challenge. **(E)** Survival curves from the re-challenge. Neutralizing antibody titers (PRNT_50_) against **(F)** rNiV and **(G)** rNiV_GhV-RBP_ were determined from sera collected PRE and POST re-challenge, respectively. Floating bar plots indicate the minimum and maximum values, with the horizontal line representing the mean.

Animals challenged with WT rNiV exhibited significant weight loss **(Figure 6B)** with all animals succumbing to infection regardless of challenge dose **(Figure 6C)**. Conversely, animals challenged with rNiV_GhV-RBP_ experienced no development of clinical signs of disease, including weight loss (**Figure 6B)** and no mortality **(Figure 6C)**. Re-challenge of the rNiV_GhV-RBP_ infected survivors with WT rNiV (dosage back-titrated to be 1.13 x 10^5^ PFU) resulted in weight loss and mortality for 1/4 animals from each challenge group; however, the remaining 3/4 animals exhibited no appreciable weight loss nor clinical signs of disease throughout the re-challenge course **(Figure 6D and 6E)**. Serum collected from survivors at 21 DPI (PRE rechallenge) and at 21 days post rechallenge with rNiV (POST rechallenge) was analyzed for neutralizing titers against rNiV and rNiV_GhV-RBP_. Following initial rNiV_GhV-RBP_ infection, serum collected 21 DPI demonstrated some neutralizing activity against rNiV, especially in the higher challenge dose. However, post rechallenge serum demonstrated an appreciable increase in neutralizing antibody titers, regardless of initial challenge dose **(Figure 6F)**. By contrast, pre rechallenge serum exhibited neutralizing activity against rNiV_GhV-RBP_, with animals in the 1×10^5^ PFU challenge group exhibiting higher overall neutralizing titers. Rechallenge with rNiV was sufficient to further boost neutralizing titers against the chimeric virus (**Figure 6G**).

Serological analysis of hamsters initially challenged with 10^5^ PFU of rNiV_GhV-RBP_ revealed the development of neutralizing antibodies against NiV and HeV by 21 DPI (**Figure S10A).** This early seroconversion likely conferred protection during the subsequent rechallenge with wild-type rNiV. Indeed, sera collected from survivors after the rechallenge exhibited a robust, broadly neutralizing response against NiV, HeV, and GhV pseudoviruses (**Figure S10B**). To evaluate the target of this polyclonal response, flow cytometry analysis of surface antibody binding confirmed recognition of NiV-F, HeV-F, and GhV-RBP in post-rechallenge sera (**Figure S10C**). Antibodies targeting NiV-F were cross-reactive only to HeV, although GhV-RBP were not cross-reactive to prototypical HNV (**Figure S10C**). Independent polyclonal responses against both HNV-F and RBP were thus capable of increasing the neutralization range against prototypical and African HNV. Because the first exposure presented GhV-RBP (in the context of NiV-F), it is most likely that anti-NiV-F antibodies were primarily responsible for protection against rNiV re-challenge.

### A panel of antivirals demonstrate efficacy against rGhV_M-GS*_

Because African bat HNVs have not been isolated in culture, their susceptibility to broad-spectrum antiviral compounds is largely unknown. We evaluated the efficacy of several broad-spectrum antivirals using inhibition assays against rGhV_M-GS*_ and rNiV in CHO-B2 cells. EIDD-2749 (4’-Fluorouridine) potently inhibited both rNiV and rGhV_M-GS*_ replication, yielding sub-micromolar IC_50_ values (0.05 and 0.06 µM, respectively) **(Figure 7A)**. Recapitulating our prior findings with GhV TC-tr minigenomes^14^, GHP-88309, an allosteric inhibitor of paramyxovirus vRdRp, was effective against rGhV_M-GS*_ (IC_50_ of 1.59 µM), but showed little if any inhibition of rNiV **(Figure 7B)**. Remdesivir inhibited both rNiV and rGhV_M-GS*_ with IC_50_ of 0.21 and 0.43 µM, respectively **(Figure 7C)**. EIDD-1931 (the active form of molnupiravir) inhibited both rNiV (IC_50_ = 3.92 µM) and rGhV_M-GS*_ (IC50 = 6.47 µM) **(Figure 7D)**. Favipiravir (T-705) demonstrated efficacy against both rNiV (IC50 = 33.14 µM) and rGhV_M-GS*_ (IC50 = 37.69 µM) (**Figure 7E**). Furthermore, EIDD-2749 **(Figure 7F)** and GHP-88309 **(Figure 7G)** demonstrated comparable efficacy against rNiV and rGhV_M-GS*_ in more physiologically relevant primary human astrocytes. A summary of all IC₅₀ values derived in this study are presented in **Fig. 7H**.

**Figure 7.**
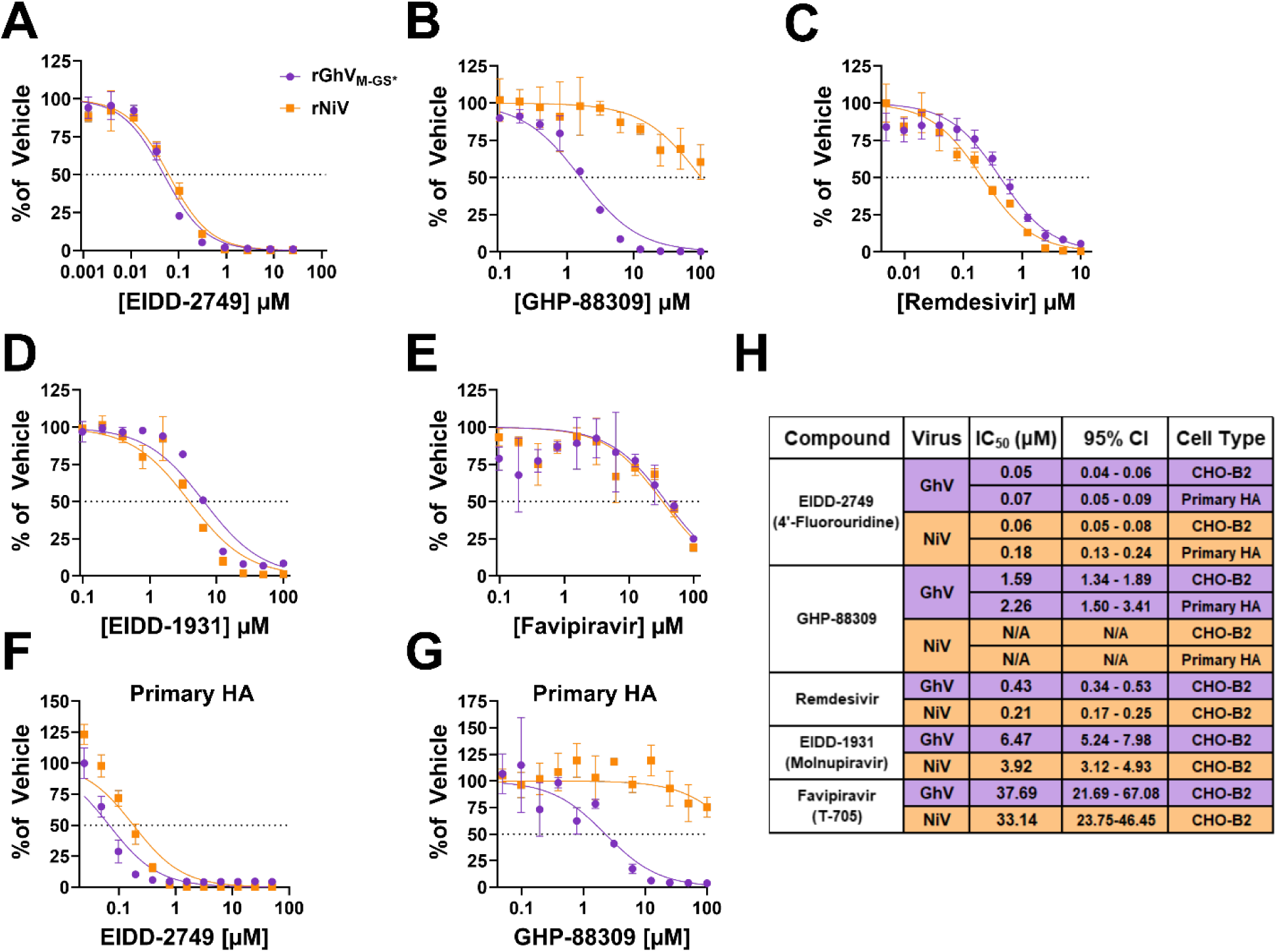
Antiviral activity of small molecule inhibitors against rGhV_M-GS*_ and rNiV. Dose-response curves are shown for various small-molecule inhibitors against rGhV_M-GS*_ (purple) and rNiV (orange). CHO-B2 cells were infected at an MOI of 0.05 and treated with **(A)** EIDD-2749, **(B)** GHP-88309, **(C)** Remdesivir, **(D)** EIDD-1931 (Molnupiravir active form), or **(E)** Favipiravir (T-705). Additional dose response curves were measured in primary human astrocytes for **(F)** EIDD-2749 and **(G)** GHP-88309 under the same MOI and assay conditions. **(H)** Summary of half-maximal inhibitory concentrations (IC_50_) derived from the dose response curves in (A-G). Each row indicates the respective compound, virus, calculated IC_50_, corresponding 95% confidence interval, and the cell type in which the assay was performed. IC_50_ values were determined in GraphPad Prism by nonlinear regression of [inhibitor] vs. normalized response. All experiments were conducted in biological triplicate. Error bars depict standard deviation.

## Discussion

### *De novo* rescue of full-length, replication-competent GhV

Difficulties in isolating viruses from environmental samples have hindered our understanding of African bat HNV biology, pathogenesis, and potential zoonotic risk. In this study, we recovered an infectious clone of GhV M74a *de novo* through reverse genetics approaches. Our prior minigenome studies, which identified a functional substitute for the unmapped genomic promoter of GhV, were crucial for recovering this infectious clone^14^. However, while encoding a functional genomic promoter into the GhV M74a sequence was sufficient to promote replication and transcription of the viral genome, the rescued virus failed to amplify efficiently beyond low-level, cell-to-cell spread. These observations suggested that there may be some defect in GhV particle assembly; in agreement with this, we identified that the GS sequence regulating matrix gene expression in GhV deviates from an otherwise conserved HNV consensus motif, and was functionally impaired in minigenome assay. Consistent with our rescue phenotype, insufficient levels of matrix protein would impede virion production and assembly, constraining virus to predominantly rely upon cell-to-cell spread for amplification. To alleviate this putative impairment, the consensus sequence was restored in the M-GS (to M-GS*) and was encoded in the full-length rGhV reverse genetics system. This virus, rGhV_M-GS*_ could be rescued and readily amplified in culture and demonstrated clear instances of both cell-to-cell and particle spread.

### Ability of GhV to antagonize human innate immune responses

The V and W proteins of NiV have been experimentally demonstrated as critical drivers of pathogenicity in animal models, and their counterparts in GhV may likewise influence the course of disease during infection^42,43^. Transcriptional profiling of rGhV_M-GS*_ demonstrated that the GhV-P gene undergoes co-transcriptional editing during GhV infection. However, editing of the GhV-P gene appears to be less frequent than is observed for NiV and HeV^23,28,29^. GhV-V inhibited MDA5-mediated signaling just as potently as NiV-V in dual luciferase assay, consistent with modeling in AlphaFold3 **(Figure 2B)**. Furthermore, the P, V, and W proteins of GhV were likewise capable of inhibiting IFN-β mediated signaling in human cells **(Figure 2C)**. These findings confirm that GhV encodes and expresses functional antagonists of human antiviral signaling; however, the reduced frequency of P gene editing (and consequently lower levels of V and W transcripts) may limit the overall potency of this antagonism during infection.

While P, V, and W from NiV have a well-established mechanism of antagonizing IFN-β signaling through the binding of STAT1 and STAT2^44–46^, our co-immunoprecipitation results suggest that antagonism of IFN-β signaling by GhV P, V, and W occurs independently of STAT1 and STAT2 binding. In agreement with this, sequence alignment of the NiV-P and GhV-P proteins demonstrates that the STAT1 binding motif in NiV-P is poorly conserved in GhV-P. Because P, V, and W from GhV are all capable of antagonism, we strongly suspect that their shared N-terminus encodes the domain or motifs responsible for antagonism. Further work is needed to better elucidate the unique mechanism behind GhV antagonism of IFN-β signaling.

### Attenuation of GhV in primary human and porcine cells

Replication of GhV is characteristically delayed and attenuated relative to NiV, with this phenotype consistently observed in both full-length virus and minigenome systems. In primary human astrocytes, GhV replication was detectable yet modest; however, infection was completely restricted in HUVECs and HBMECs. Because NiV infection of the brain microvasculature is thought to facilitate entry into the central nervous system, our findings suggest that GhV may be unable to effectively cross the blood-brain barrier. This may offer a potential barrier to neuroinvasion and pathogenicity during GhV infection. However, primary porcine kidney epithelial cells supported moderate replication of rGhV_M-GS*_, indicating that pigs may be a compatible host. Because domestic pigs served as amplifying hosts during the initial outbreak of NiV in Malaysia and Singapore^47,48^, our findings raise concerns that pigs may provide a similar niche for GhV and its close relatives. Whether this concern applies to other divergent African bat HNVs remains to be seen. Concerningly, agricultural serosurveys conducted in Ghana, Nigeria, and Uganda have demonstrated instances of seropositivity towards NiV in pigs, implying that spillover events may already be occuring^49–51^.

### Differential infection of GhV and NiV in bat cells

Our infection studies in bat cell lines suggests that GhV is adapted to its natural hosts and may rely upon bat-specific cellular factors for optimal replication. Notably, *E. helvum* derived ZFB<u>S13-75A</u> cells supported robust rGhV_M-GS*_ replication without the characteristic kinetics delay observed in non-chiropteran cells **(Figure 4C)**. Similarly, *E. gambianus* ZFBK11-97 cells exhibited robust replication and pronounced GhV CPE **(Figure 4F and S8F)**; this is consistent with previous studies demonstrating that GhV fusogenicity may be relatively restricted to specific chiropteran cell lines^52–54^. Given that *E. gambianus* is sympatric with *E. helvum*, the former may be naturally exposed to GhV or related African HNVs. However, serological data from Ghana show low reactivity to NiV antigen in *E. gambianus* (1%; 1/89), in sharp contrast to high seroprevalence in *E. helvum* (39%; 23/59)^7^, suggesting species-specific differences in exposure or susceptibility.

Other bat species investigated in this study, including *P. dasymallus*, *R. ferrumequinum*, and *R. leschenaultii*, do not share an overlapping geographic range with *E. helvum*, and are unlikely to encounter GhV in nature. Nonetheless, our findings offer insights into the relative permissivity of these bats to divergent HNVs and provide a framework for evaluating potential susceptibility across bat genera. The overall infectivity profile of GhV in bat cells correlated with relative EFNB2 transcript levels, with cells that express higher levels of EFNB2 (ZFB<u>S13-75A</u>, BKT1, and ZFB<u>K11-97</u>) generally supporting better GhV replication. By contrast, NiV infection did not exhibit such a relationship, suggesting that GhV entry is more sensitive to receptor abundance than NiV. Prior structural studies offer a mechanistic explanation for this observation, where GhV-RBP is observed to engage EFNB2 in a similar manner to NiV-RBP, but lacks a secondary interaction site present in NiV-RBP that enhances receptor affinity and stabilizes the virus-receptor complex^22^. These data support a threshold model wherein NiV can efficiently infect cells with relatively low EFNB2 expression, while GhV requires higher receptor density to overcome its weaker binding and less efficient fusion activation.

Notably, *R. ferrumequinum* BKT1 and *R. leschenaultii* DemKT1 cells supported overall greater replication of rGhV_M-GS*_ than rNiV, suggesting a species-specific restriction of NiV. The G-H loop of EFNB2, which interacts with NiV-RBP and GhV-RBP for binding, is absolutely conserved across the bat species in this study **(Figure S8J-K)**, suggesting the restriction is not at the level of entry. Previous studies have demonstrated that experimentally-challenged *Rousettus aegyptiacus* bats do not support productive NiV or CedV infection *in vivo*^55,56^. Furthermore, NiV does not productively replicate in primary *R. aegyptiacus* cells^55^, suggesting that the block we observe in *R. leschenaultii* cells may reflect a broader, genus-level restriction mechanism. However, there is serological and metagenomic evidence to suggest that *Roussettus spp.* bats in Africa do harbor divergent HNVs^34,57^, and a recent study identified two new HNV species through metagenomic analysis of *R. leschenaultii* bats in China^58^. This raises the possibility that particular HNV clades may have evolved to be more compatible with *Rousettus* biology than other species, such as NiV. For *Rhinolophus* spp., experimental infection with NiV has not been reported, and there is limited evidence of natural infection in this genus, leaving their compatibility with HNVs largely unexplored^59^. Future studies will aim to identify the mechanisms of post-entry restriction specific to NiV, but not GhV, in these cells.

### GhV and an EFNB3-blind NiV are non-pathogenic in Syrian golden hamsters

Our *in vivo* work demonstrates that, despite driving seroconversion, GhV is attenuated in Syrian golden hamster model. This attenuation may stem from a combination of its limited receptor usage and slower replication kinetics. These findings provide the first experimental evidence to suggest that GhV may be non-pathogenic; however, additional studies in other animal models are required to better determine if these results are generalizable. EFNB3 receptor usage has been speculated to play a role in the neurotropism and neurovirulence associated with NiV and HeV infection^17^. In support of this, *in vivo* work has recently demonstrated that Cedar virus, a non-pathogenic HNV which uses EFNB2 but not EFNB3, is unable to infect the central nervous system of experimentally-infected IFNAR-KO mice^60^. However, CedV also lacks the V and W accessory proteins, which in other HNVs function as potent antagonists of the host antiviral response; in NiV, experimental knockout of the V protein was sufficient to attenuate disease in hamsters and ferrets^42,43^. As a result, it is difficult to disentangle the relative contributions of receptor usage and innate immune evasion in shaping CedV’s restricted tropism and non-pathogenic phenotype in hosts^61–64^.

Because our rNiV_GhV-RBP_ chimera maintains the authentic NiV replicative machinery and full repertoire of potent NiV innate immune antagonists, our findings strongly implicate EFNB3 usage as a major determinant of HNV pathogenesis. More broadly, these results highlight that receptor usage and innate immune antagonism are likely to act in tandem to govern host tropism and immune evasion, ultimately resulting in severe disease outcome. Furthermore, the ability of the rNiV_GhV-RBP_ chimeric virus to elicit atypically broad antibody responses and protection against WT NiV in hamsters suggests that presentation of heterotypic envelope proteins could be valuable for developing vaccine approaches to elicit polyclonal responses with the ability to neutralize divergent HNVs.

### GhV susceptibility to broad-spectrum antivirals

Our profiling of GhV and NiV susceptibility to small-molecule inhibitors aligns with previous studies demonstrating that EIDD-2749, Remdesivir, and Favipiravir are active against NiV^65–68^. Validating previous work in our rGhV TC-tr minigenome, GHP-88309 was demonstrated to be effective against rGhV_M-GS*_ but not rNiV^14^. The resistance of NiV to GHP-88309 reflects structural differences in the viral polymerase that prevent effective binding of the compound, consistent with prior studies^68–70^. To our knowledge, this study also represents the first report of inhibition of HNV replication by EIDD-1931, the active form of molnupiravir. Future studies are warranted to validate the efficacy of molnupiravir *in vivo,* and to better determine if this drug, which has been employed in the clinic, may be appropriately repurposed to treat henipaviral infections.

## Limitations of the study

Despite the insights gained from this study, we acknowledge certain limitations that may influence the interpretation and generalizability of our findings. First and foremost, we acknowledge that the infectious clone of GhV M74a employed in this study (rGhV_M-GS*_) likely diverges from authentic, circulating GhV in certain respects. Specifically, the HeV derived Ldr28 sequence utilized in lieu of the unmapped genomic promoter may not reflect the authentic, unknown genomic promoter of GhV, although it was stably maintained during amplification of our viral stocks as confirmed by RNA sequencing.

Furthermore, with only a single reference sequence and a dearth of environmental isolates, we cannot definitively say if the non-canonical M-GS sequence reported in GhV M74a is representative of circulating quasispecies or if it is an artifact of sequencing. Similar viral engineering was necessary to achieve the rescue of Lloviu virus from its reference sequence, which likewise lacked genomic ends and necessary cis-acting elements due to sequencing artifacts^71^. These difficulties highlight the challenges associated with recovering virus from sequence alone. Nevertheless, the CDS of all GhV M74a genes remain unchanged from the reference sequence in GhV_M-GS*_, and we argue that the observed biological phenomena (receptor usage, pathogenesis, and post-entry compatibility) are faithful to authentic GhV biology.

## Concluding remarks

In summary, this study provides the first experimental characterization of a full-length, replication-competent African bat henipavirus, and yield valuable insights into its biology, host range, and zoonotic potential. Comparative infection profiling of NiV and GhV across a variety of cell lines revealed that while GhV exhibits limited replication with delayed kinetics in non-chiropteran cells, GhV replication in bat cells is more robust. In agreement with its attenuated phenotype in non-chiropteran species, infection of Syrian golden hamsters with GhV did not result in disease. Furthermore, we confirm EFNB3 usage as a major determinant of HNV pathogenicity by constructing a chimeric, EFNB3-blind NiV that was completely attenuated *in vivo*. Collectively, our findings suggest that GhV may require further adaptation to achieve efficient replication in non-chiropteran hosts, and even in the event of spillover, its inability to utilize EFNB3 may result in subclinical infection. Nevertheless, our observation that primary human and porcine cells support even low-level GhV replication highlights the need for continued research and surveillance efforts to prevent, identify, and mitigate future spillover events of African bat HNVs.

## Resource Availability

Sequences of the GhV reverse genetics plasmids have been deposited in GenBank under accession numbers PX138233–PX138236. Whole-genome sequencing of rGhV stocks has been deposited in NCBI BioProject PRJNA1284405. Long-read direct RNA sequencing of GhV-infected cells has been deposited in NCBI BioProject PRJNA1307648. All numerical data underlying figures in this manuscript will be deposited in Mendeley Data and made publicly available upon publication.

## Acknowledgements.

This work was supported in part by NIH grants R01 AI185102 (B.L.) and U19 AI171403 (B.L., A.N.F., A.L.G.). B.L acknowledges support by the Bill and Melinda Gates Foundation PAD Henipavirus Initiative (INV-048877). G.D.H. acknowledges support by the National Science Foundation Graduate Research Fellowship Program (Grant No. 1842169). Any opinions, findings, and conclusions or recommendations expressed in this material are those of the authors and do not necessarily reflect the views of the National Science Foundation.

## Star Methods

### Maintenance of cell lines

BSRT7 (CVCL_RW96), Vero CCL81 (CVCL_0059), HEK-293T (CVCL_0063), BKT1 (CVCL_YZ67), DemKT1 (CVCL_YZ68), ZFBK13-75, and EidNi/41.2 (CVCL_RX13) cells were maintained in Dulbecco’s modified Eagle medium (DMEM) supplemented with 10% fetal bovine serum (FBS, GeminiBio). ZFBS13-75A (CVCL_YZ77) and ZFBK11-97A (CVCL_YZ74) were maintained in RPMI 1640 media supplemented with 10% FBS. The generation of CHO pgsA-745 cells overexpressing EFNB2 or EFNB3 (CHO-B2 and CHO-B3) have been previously described^17^, and were maintained in DMEM/F-12 media supplemented with 10% FBS. Primary human cells were sourced from ScienCell Research laboratories, including human astrocytes (catalog #1800), human brain microvascular endothelial cells (catalog #1000), and human umbilical vein endothelial cells (catalog #8000). All primary cells were maintained at low passage, with HUVECs and HBMECs maintained in endothelial cell medium with kit (ScienCell catalog #1001) and HAs maintained in astrocyte medium with kit (catalog #1801). Normal human bronchial epithelial cells (Lonza, catalog #CC-2540) were grown in BEGM bronchial epithelial cell growth media with kit (Lonza, catalog #CC-3170). Small airway epithelial cells (Lonza, catalog #CC-2547) were maintained in SAGM small airway epithelial cell growth media with kit (Lonza, catalog #CC-3118). Primary porcine kidney epithelial cells (CellBiologics #P-6034) were maintained in complete epithelial cell medium with kit (CellBiologics #M6621). All cells were grown at 37°C in 5% CO_2_.

### Design and cloning of reverse genetics plasmids

The full antigenomic sequence of rGhV strain Eid_hel/GH-M74a/GHA/2009 (NCBI Reference Sequence NC_025256.1) was fully synthesized (Bio Basic Inc.) and assembled into the pEMC vector. The non-coding region (NCR) between the N and P genes was duplicated to accommodate insertion of the GLuc-P2A-eGFP reporter, and there is an artificial NotI restriction site immediately downstream of the reporter gene’s stop codon. The HeV Ldr28 sequence (the terminal 3’-most 28 nucleotides of the HeV 3’ leader sequence) was inserted upstream of the reported GhV M74a sequence as previously described in minigenome^14^. The viral antigenomic sequence is flanked on its 5’ terminal end by a sequence-specific hammerhead ribozyme element (5’-AGATCGGTCTGATGAGTCCGTGAGGACGAAACGGAGTCTAGACTCCGTC-3’) and on its 3’ terminal end by the HDV ribozyme, as has been detailed for our paramyxovirus reverse genetics systems^38,72,73^. An optimal T7 promoter sequence lies upstream of the 5’ hammerhead ribozyme sequence, and a T7 terminator sequence lies downstream of the 3’ HDV ribozyme sequence. To restore the consensus motif in the M-GS sequence (M-GS*), a combination of restriction digest, PCR, and InFusion cloning were employed. Whole-plasmid sequencing confirmed that rGhV and rGhV_M-GS*_ antigenome plasmids (beyond reporter gene integration and modification of the M-GS) matched the GhV M74a reference sequence with the exception of a single-nucleotide change in the 3’ UTR of the GhV-N gene, in which there was a C1851T mutation.

The accessory plasmids encoding GhV-N, -P, and -L, and the rGhV TC-tr minigenome were generated as described previously^14^. The native sequence of GhV-N, -P, and -L genes, respectively, were encoded into the pTM1 vector downstream of the EMCV IRES, between the SpeI and XhoI restriction sites, as previously described for NiV and HeV accessory plasmid construction^74^. The ORF of the C protein was silenced in the GhV-P accessory plasmid by silently mutating its start codon. For some constructs, the N-GS in the rGhV TC-tr minigenome was replaced with the M-GS and M-GS* sequences, respectively. Sequences of the GhV reverse genetics plasmids have been deposited in NCBI GenBank (Accession numbers PX138233-PX138236).

The accessory plasmids for NiV (NiV-N, -P, and -L) were constructed and described previously^38^. The rNiV (Malaysia strain UMMC1) encoding eGFP and *Gaussia* luciferase was rescued as previously described^11,38,39^. The rNiV_GhV-RBP_ chimera was constructed by replacing the CDS of the NiV-RBP gene with the CDS of the GhV-RBP gene. Cloning was achieved in all cases by a combination of restriction enzyme digest and InFusion-mediated fragment assembly (TakarBio). All restriction enzymes were purchased from New England Biolabs, and all primers employed were synthesized by Millipore Sigma.

### TC-tr minigenome assays

BSRT7 cells were seeded one day prior to transfection in 24-well format, to achieve 70-80% confluence on the day of transfection. Twenty-four hours later, cells were co-transfected with plasmids encoding respective the rHNV TC-tr minigenome, codon-optimized T7 polymerase, and HNV-N, -P, and -L as previously described^14^. All transfections used Lipofectamine LTX and PLUS according to the manufacturer recommendations. 1.39 micrograms of respective TC-tr minigenome, 0.40 micrograms of codon-optimized T7 polymerase, 0.49 micrograms of HNV-N, 0.32 micrograms of HNV-P, 0.16 micrograms of HNV-L, 1.27 microliters of PLUS, and 2.04 microliters of LTX per reaction constituted each minigenome reaction. For all transfections, mastermixes were prepared which contained the TC-tr minigenome, HNV-N, HNV-P, and codon-optimized T7; these master mixes were then aliquoted into equal parts before adding plasmid encoding HNV-L or plasmid encoding GFP. Plasmid encoding GFP was used in lieu of HNV-L as a control to determine background signal nonspecific to the viral replicase. At 24 HPT, media was replaced on plates. Quantification of mCherry events and nanoluciferase assays were conducted at 48-72 hours post-transfection, depending on the experiment.

To measure nanoluciferase activity, TC-tr minigenome cells were lysed and processed using the Nano-Glo® HiBiT Lytic Detection System (Promega Corporation), according to the manufacturer recommendations. Cell lysate was transferred to white 96-well plates and RLUs were measured on a Cytation 3 plate reader (BioTek) using a gain of 125 and an integration time of 1.0s. Raw RLUs were then normalized to the average RLUs of the no-L (GFP) control to determine the vRdRp-dependent signal above background. A Celigo Imaging Cytometer (Nexcelom) was used to image the entire plate of TC-tr minigenome cells in the red channel. The total number of mCherry positive events were quantified using Celigo software analysis of the red channel.

### Rescue and amplification of full-length rGhV

350,000 BSRT7 cells per well were seeded into 6-well plates. Twenty-four hours later, when cells were approximately 70% confluent, the media on each plate was exchanged for 1.5mL of fresh DMEM + 10% FBS containing 1x Nucleic Acid Transfection Enhancer (Invivogen). Transfection reactions were prepared by diluting 4.0 micrograms of plasmid encoding codon-optimized T7 polymerase, 5.0 micrograms of pTM1_T7-GhV-N, 3.2 micrograms of pTM1_T7-GhV-P in 200 microliters of OptiMEM. 21 microliters of Mirus TransIT-LT1 were then added to the transfection reaction and gently mixed. After a 30 minute incubation at room temperature, transfection complexes were added dropwise to BSRT7 cells. At 24 hours post-transfection (HPT), media was exchanged on every well with 1.5mL of fresh DMEM + 10% FBS containing 1x Nucleic Acid Transfection Enhancer (Invivogen). The cells were then moved into high containment (Galveston National Laboratory BSL-4, University of Texas Medical Branch) for transfection of the antigenome. The rGhV antigenomic plasmid was diluted in 200 microliters of OptiMEM before adding 21 microliters of Mirus TransIT-LT1 to the reaction. After gently mixing, transfection complexes were incubated at room temperature for 30 minutes prior to being added dropwise to the replicase-transfected cells.

Rescue cells were monitored daily for signs of CPE and reporter gene expression. To maintain cell viability, media was exchanged every other day (DMEM containing 10% FBS and 1x Nucleic Acid Transfection Enhancer). Upon observation of obvious secondary spread, supernatant was harvested daily with replacement and frozen at - 80°C. Stocks of rGhV_M-GS*_ were made by infecting Vero CCL81 and CHO-B2 cells, respectively, with P0 supernatant. Upon observation of widespread (>90%) virus amplification in flask, as determined by CPE and GFP expression, infected flasks were freeze-thawed once and supernatant was harvested, clarified by brief centrifugation, and frozen at −80°C.

### Rescue and amplification of rNiV and rNiV_GhV-RBP_

Rescue and amplification of rNiV (Malaysia strain) constructs was conducted as previously described^11,38,39^. BSRT7 cells were seeded at a density of 400,000 cells per well into six-well plates. A day later, when cells had reached 80% confluence, transfection reactions were prepared by diluting 7.0 micrograms of rNiV or rNiV_GhV-RBP_ antigenomic plasmid, 2.0 micrograms of codon optimized T7 polymerase, 2.5 micrograms of pTM1_T7-NiV-N, 1.6 micrograms of pTM1_T7-NiV-P, and 0.8 micrograms of pTM1_T7-NiV-L in 200 microliters of OptiMEM. Cells and plasmid mixes were then moved to the high containment, where 21 microliters of Mirus TransIT-LT1 was added to the transfection reaction and gently mixed. After a 30 minute incubation at room temperature, transfection complexes were added dropwise to the BSRT7 cells. Rescue cells were monitored daily for signs of infection (CPE and reporter gene signal). Upon observation of obvious secondary spread, media was harvested with replacement and stored at −80°C or used to infect Vero CCL81 cells. Upon widespread virus amplification in the flask, as determined by CPE and GFP expression, the flask was freeze-thawed once and supernatant was harvested, clarified by brief centrifugation, and frozen at −80°C.

### Virus work and biosafety

All work with full-length, infectious rNiV and rGhV was conducted in class II BSC at the Galveston National Laboratory BSL-4, and at The Robert E. Shope, MD, Laboratory BSL-4 at the University of Texas Medical Branch (UTMB). All samples requiring removal from high containment were rendered inactivated/non-infectious using institutionally-approved protocols, and samples were rigorously validated to be non-infectious prior to removal.

### Virus titration

Stocks and infectious samples of rNiV and rNiV_GhV-RBP_ were titrated by plaque assay. 250,000 cells/well of Vero CCL81 cells were seeded in 12-well format. Twenty-four hours later, virus was serially diluted in DMEM + 10% FBS and cells were infected in low volume (100 microliters) for 1 hour at 37°C. Following this 1 hour incubation, semi-solid overlay was added onto each well (1x MEM + 0.5% methylcellulose + 2%FBS) and plates were incubated at 37°C until assay endpoint. Plates were fixed in crystal violet buffered in 10% formalin and read at 72 HPI for rNiV or 5 DPI for rNiV_GhV-RBP_.

Because rGhV_M-GS*_ did not generate sufficient CPE for plaque assay, we titrated virus on highly permissive CHO-B2 cells via TCID_50_. 96-well plates were seeded with 15,000 cells/well of CHO-B2 cells. The next day, virus was serially-diluted in DMEM/F-12 + 10% FBS and incubated on CHO-B2 cells overnight at 37°C. At 24 HPI, media was replaced on all wells to minimize residual *Gaussia* luciferase signal from the inoculum. At 72HPI, supernatant was harvest for *Gaussia* luciferase assay. Infected wells were determined by RLU signal and titer (TCID_50_/mL) was determined using the Reed-Muench method^75^. All TCID_50_ assays were conducted using 4 to 6 technical replicates per sample.

### Virus growth kinetics

To determine the growth kinetics of rGhV_M-GS*_ and rNiV, 200,000 cells/well of CHO-B2 cells were seeded into 12-well format. The following day, cells were infected in triplicate, using low-volume inoculum (100 microliters), with either rGhV_M-GS*_ or rNiV at an MOI of 0.05 for 1 hour at 37°C. Plates were gently rocked every 15 minutes to prevent drying and to ensure even distribution of the inoculum. The titer of rGhV_M-GS*_ was converted to PFU/mL by multiplying the TCID_50_/mL titer by 0.7. After the 1 hour incubation, viral inoculum was removed and cells were gently washed with 1mL of DPBS to remove residual GLuc and unbound virus. The DPBS was removed, and 1.5 mL of DMEM:F12 + 10% FBS was added to the wells. We then harvested and froze 0.5 mL of supernatant at −80°C, which served as the “0 HPI” sample. After this, 0.5 mL of supernatant was harvested every 24 hours with replacement and froze at −80°C. All timecourse samples were subsequently thawed for use in virus titration and *Gaussia* luciferase assays.

### Interferon stimulated response reporter gene assays

The NiV and GhV-P, -V, and -W genes were subcloned from the pTM1_T7-NiV-P and pTM1-T7-GhV-P plasmids, with PCR used to append a triple FLAG tag onto the N-terminus of each gene (CloneAmp, TakaraBio). InFusion Cloning (TakaraBio) was used to assemble the respective gene products into pCAGGs expression vector.

For luciferase assays, HEK-293T cells were transfected with: (1) the construct encoding the respective P, V, W, or EV sequence, (2) a plasmid containing Firefly luciferase gene downstream of an IFN-stimulated response element (pISRE-TA-Luc), and (3) a construct encoding the Renilla luciferase protein (pRL-Luc). Transfections were performed using lipofectamine LTX and PLUS (Invitrogen).

To test antagonism of MDA5-mediated signaling, plasmid encoding FLAG-tagged MDA5 under the CAGG promoter was co-transfected into cells in parallel with the plasmids above. At 24 HPT, cells were lysed in 1x passive lysis buffer (Promega). To test antagonism of IFN-β mediated signaling, at 24 HPT, cells were treated with 1000 units of human IFN-β and incubated at 37°C for 6 hours. After the 6 hour incubation period, cells were lysed in 1x passive lysis buffer. Lysates were employed in SDS-PAGE and Dual-Glo® Luciferase Assay System (Promega). Firefly luciferase values were normalized to the Renilla control to account for differences in transfection.

### Western blotting

Protein content in lysates were quantified using Bradford Assay, and equal amounts of protein were combined with 1x Laemelli buffer with DTT and boiled for 5 minutes 95°C before being transferred onto ice. Samples were loaded onto pre-cast, 4-20% mini-PROTEAN polyacrylamide gels, and proteins were resolved via electrophoresis in the Mini-PROTEAN Tetra cell (BioRad). Proteins were transferred to PVDF membranes using the Trans-Blot Turbo transfer system (BioRad). Membranes were blocked with phosphate-buffered saline blocking buffer (LI-COR; 927–700001) and then probed with the indicated antibodies. Antibodies against FLAG (Sigma-Aldrich, F3165), GAPDH (Cell signaling, 2118), STAT1 (Cell signaling, D1K9Y), and STAT2 (Cell signaling, D9J7L) were employed. Secondary antibodies conjugated to Alexa Fluor 647 or Alexa Fluor 546 were used to detect respective primary antibodies. For western blotting of samples from the GhV TC-tr minigenome transfections, proteins were transferred to nitrocellulose membranes and probed for HiBiT using the Nano-Glo HiBiT blotting system (Promega) according to manufacturer instructions. All blots were imaged using a ChemiDoc MP imaging system (Bio-Rad).

### Co-immunoprecipitation assays

HEK-293T cells were transfected with 200ng of respective FLAG-tagged viral protein or EV using Lipofectamine LTX and PLUS. At 24 HPT, cells were treated with 1000 U/mL IFN-β for 6 hours, lysed, and lysates were subjected to immunoprecipitation using anti-FLAG beads (Sigma-Aldrich, A2220). Western blots were performed as described for other assays, with IP eluates and whole-cell lysates probed to detect FLAG (viral proteins), STAT1, and STAT2.

### Timecourse for primary cells

To determine the growth kinetics of rGhV_M-GS*_ and rNiV in primary human cells, 100,000 cells/well of HAs, HUVECs, or HBMECs were seeded into poly-L-lysine coated 24-well format. The following day, cells were infected in triplicate, using low-volume inoculum (100 microliters), with either rGhV_M-GS*_ or rNiV at an MOI of 0.1 for 1 hour at 37°C. Plates were gently rocked every 15 minutes to prevent drying and to ensure even distribution of the inoculum. The titer of rGhV_M-GS*_ was converted to PFU/mL by multiplying the TCID_50_/mL titer by 0.7. After the 1 hour incubation, viral inoculum was removed and cells were gently washed with 0.5 mL of DPBS to remove residual GLuc and unbound virus. The DPBS was removed, and 0.6 mL of respective cell-specific media was added to the wells. We then harvested and froze 0.1 mL of supernatant at - 80°C, which served as the “0 HPI” sample. After this, 20 microliters of supernatant was harvested every 24 hours with replacement and froze at −80°C. All timecourse samples were subsequently thawed for use in virus titration (0HPI vs endpoint) and *Gaussia* luciferase assays.

For timecourse studies with NHBE and SAEC, 25,000 cells/well were seeded into 96-well format. For PPKE, 15,000 cells/well were seeded into 96-well format. Cells were then infected with rNiV or rGhV_M-GS*_ at an MOI of 0.1 for 1 hour at 37°C. Cells were then rinsed with 0.1 mL of DPBS, and 120 microliters of respective media was added to each well. 20 microliters of media was then harvested and frozen at −80°C, serving as the “0 HPI” timepoint. After this, 20 microliters of supernatant was harvested every 24 hours with replacement and froze at −80°C. All timecourse samples were subsequently thawed for use in *Gaussia* luciferase assays.

### Timecourse for immortalized cells

For timecourse studies using immortalized cells (CHO-B2, CHO-B3, ZFBS13-75A, ZFBK13-75, ZFBK11-97, FBKT1, BKT1, and DemKT1) 15,000 cells/well were seeded into 96-well format. A day later, cells were then infected with rNiV or rGhVM-GS* at an MOI of 0.05, as titered on CHO-B2 cells, for 1 hour at 37°C. After the 1 hour infection, cells were rinsed with 0.1 mL of DPBS (to remove residual *Gaussia* luciferase from the virus stock), and 120 microliters of respective media was added to each well. 20 microliters of media were then harvested and frozen at −80°C, serving as the “0 HPI” timepoint. After this, 20 microliters of supernatant was harvested with replacement at denoted timepoints and frozen at −80°C. All timecourse samples were subsequently thawed at the same time for use in *Gaussia* luciferase assays.

### *Gaussia* luciferase assays

*Gaussia* luciferase assays were carried out using the Pierce Gaussia Luciferase Glow Assay kit (Thermo Fisher Scientific), according to manufacturer instructions. In general, 20 microliters of sample were gently mixed with substrate diluted in *Gaussia* glow assay buffer in opaque, black-bottom 96-well plates. After a ten minute incubation at room temperature, plates were read for luminescence on a Cytation5 (Biotek) plate reader.

For timecourse studies of rNiV and rGhV, 20 microliters of supernatant were collected with replacement at each time point and frozen at −80°C. Once all timepoints had been collected, samples were thawed simultaneously and used for *Gaussia* luciferase assay as described above. The RLUs at each timepoint were then normalized to the average RLUs of each condition at 0 HPI to gauge fold-change in RLUs over time.

### *In vivo* hamster challenges

All animal studies were approved by the IACUC committee at UTMB (protocol # 1302006, and 2204021). Groups of five 5-6 week old, equal-sex Syrian golden hamsters (Envigo, Indianapolis, IN, USA) were challenged as described using respective virus diluted in volume of 100 microliters via the intraperitoneal (IP) or intranasal (IN) route. Back titration of inocula was performed on the challenge day when possible and is reported. Hamsters were monitored at least once daily until clinical symptoms onset, and up to 3 times a day during peak disease. Body weights were taken daily for the first 9-10 days, then every 3 days for the remainder of the study.

Moribund animals (determined via clinical scoring or displaying greater than 20% weight loss) were euthanized. Clinical scoring included change in breathing (labored, irregular) and a bloody nose, while criteria of a neurological involvement included aggressive behavior, any form of paralysis, ataxia, head tilt, and/or seizure. Retroorbital bleeds were carried out at 2, 4, 6, and 8 DPI. At time of euthanasia, terminal serum was first collected via cardiac puncture. For each manipulation (viral infection or biosampling), animals were anesthetized with isoflurane (Piramal, Bethlehem, PA).

### Detection and quantification of neutralizing antibody titers from serum

To determine the titer of neutralizing antibodies from animal sera against rNiV or rNiV_GhV-RBP_, plaque reduction neutralization tests (PRNTs) were performed in 12-well format. Serum was heat inactivated at 56 °C for 30 minutes prior to serial dilution in MEM with 2%FBS. Diluted sera was incubated with a dose of 40 PFU of rNiV or rNiV_GhV-RBP_ for one hour at 37°C. Vero cells were then infected with the virus:serum mix, and a standard plaque assay was conducted. PRNT_50_ was defined as the antibody dilution yielding 50% reduction in plaques.

To determine the titer of neutralizing antibodies from animal sera against rGhV_M-GS*_, luciferase reduction neutralization tests (LRNTs) were performed in 96-well format. Serum was heat inactivated at 56 °C for 30 minutes prior to serial dilution in DMEM/F-12 with 2% FBS. Diluted sera was incubated with a dose of 1500 PFU of rGhV_M-GS*_ for one hour at 37°C. CHO-B2 cells were then infected with the virus:serum mix. Media was changed at 24 HPI, and supernatant was collected at 72 HPI for *Gaussia* luciferase assay. LRNT_50_ was defined as the antibody dilution yielding a 50% reduction in RLUs relative to mock.

### Drug inhibition assays

For drug inhibition assays, 15,000 cells/well of CHO-B2 or primary human astrocyte cells were seeded in 96-well format and infected the next day with rNiV or rGhV_M-GS*_ at an MOI of 0.05. Serially-diluted compounds were added at the time of infection. At 24 HPI, the supernatant was replaced with fresh DMEM + 10% FBS containing respective compound dilutions. At 48 HPI, 20 microliters of supernatant was collected and transferred to an opaque, black-bottom plate for *Gaussia* luciferase assay. Raw RLUs were normalized to the DMSO control and multiplied by 100 to determine signal in each condition relative to vehicle. All drug inhibition assays were conducted in at least biological triplicate, and IC_50_ values were calculated in GraphPad Prism software by nonlinear regression of the [inhibitor] vs normalized response. Compounds employed in this study include EIDD-2749 (MedChemExpress, HY-146246), GHP-88309 (Sigma Aldrich, SML2997), EIDD-1931 (MedChemExpress, HY-125033), Remdesivir (MedChemExpress HY-104077), and Favipiravir (Cellagen Technology, C8705-5). All compounds were diluted according to manufacturer recommendations.

### Long read direct RNA sequencing

CHO-B2 cells were infected with rGhV_M-GS*_ at an MOI of 0.05. At 72 HPI, total RNA was collected using 0.5 mL of TRIzol reagent. Direct-zol RNA Miniprep kits were used to extract total RNA (Zymo Research). RNA was eluted in ultra-pure water and stored at −80°C. RNA samples were demonstrated to be non-infectious in CHO-B2 cells, which showed no reporter gene activity nor CPE for two passages in high containment. Total RNA was prepared for sequencing library using the Direct RNA sequencing kit (Oxford Nanopore Technologies, SQK-004), and libraries were sequenced on a RNA sequencing flow cell (FLO-MIN004RA). Basecalling was achieved using Dorado 7.6.7 in the MinKNOW 24.11.8 software, and reads were aligned to the rGhV_M-GS*_ genome using Minimap2^76^. P editing was quantified using mPileup in Samtools, selecting for reads with inserts of Gs only^77^. Raw sequencing reads have been deposited in NCBI BioProject ID PRJNA1307648.

### Sequencing of GhV stocks

Whole-genome sequencing of the four rGhV stocks was conducted using a metagenomic next-generation sequencing approach as previously described^78^. Libraries were sequenced on an Illumina NextSeq 2000 platform with 2×150bp reads. Raw reads were trimmed and quality filtered with fastp (v0.23.4)^79^. Filtered reads were then used for variant calling using the RAVA workflow (default parameters) and the plasmid map sequence as a reference (https://github.com/greninger-lab/RAVA_Pipeline/tree/2025-06-30_GhV_Griffin_et_al)^80,81^. Raw sequencing reads have been deposited in NCBI BioProject PRJNA1284405.

### RT-qPCR for EFNB2 and EFNB3

Total RNA was extracted from respective bat cell lines using TRIzol reagent and the Direct-zol RNA Miniprep kit (Zymo Research). RNA was eluted in ultra-pure water and stored at −80°C. The Primer-free LunaScript RT Master Mix Kit (New England Biolabs) was used for first-strand cDNA synthesis using gene-specific primers targeting EFNB2, EFNB3, or GAPDH. A no-RT control was employed in parallel. Following cDNA synthesis, equal volumes of cDNA were used for qPCR using the Luna Universal qPCR Master Mix kit (New England Biolabs). For qPCR, gene specific primers were utilized targeting either a 189-basepair region of the EFNB2 gene (FWD: 5’-AGGGACTCCGTGTGGAAGTA-3’ ; REV: 5’-AGAGTCCACTTTGGGGCAAAT-3’), a 169-basepair region of the EFNB3 gene (FWD: 5’-TCTCCGCTTCACCATCAAGT-3’ ; REV: 5’-TCGGAGAAGCACCTTCATGC-3’), or a 166-basepair region of the GAPDH gene (FWD: 5’-TCAAGGGCATCCTGGGCTA-3’ ; REV: 5’-ACCACCCTGTTGCTGTAGCCAA-3’). Signal was captured on a C1000 Touch Thermal Cycler (Bio-Rad), and data were exported for analysis. All qPCR reactions were conducted in at least biological duplicate. Relative gene expression was determined by first calculating the ΔCT of EFNB2 (CT_EFNB2_ - CT_GAPDH_) and the ΔCT of EFNB3 (CT_EFNB3_ - CT_GAPDH_). Then, all ΔCT values for EFNB2 or EFNB3, respectively, were normalized to ZFBK13-75 by calculating ΔΔCT(ΔCT_X_ – AVG[ΔCT_ZFBK13-75_]). Values were plotted as 2^-^_ΔΔCT._

### Microscopy

For microscopic imaging, the nuclei of cells were stained using NucBlue™ Live ReadyProbes™ Reagent (ThermoFisher) at a concentration of 2 drops/mL; cells were incubated in NucBlue-containing solution for at least 30 minutes prior to imaging. Microscopic images were collected using an Olympus IX83 microscope. All images at a given timepoint were collected using the same settings for each channel. Images were subsequently processed in ImageJ software.

### *In silico* analyses of GhV gene CDS

The CDS of all GhV M74a genes were evaluated through a combination of *in silico* approaches. The amino acid sequences of each GhV protein were aligned to their homologues from NiV, strain UMMC1 (GenBank AY029767.1), HeV strain HeV/Australia/1994/Horse18 (GenBank MN062017.1), Cedar virus strain CG1a (GenBank JQ001776.1), and Angavokely virus (GenBank ON613535.1). Alignments were conducted in MegaX using Clustal Omega, and were manually inspected. Amino acid percent identity tables were generated using Clustal Omega webserver (https://www.ebi.ac.uk/jdispatcher/msa/clustalo). Phylogenies were created in MegaX using the neighbor-joining method with 1000 bootstrap iterations^82^. Structural comparisons between GhV and NiV were conducted by either: (1) comparing resolved structures available on PDB, or (2) comparing predicted models generated using AlphaFold3. All structural analyses were conducted in UCSF ChimeraX, and RMSD was determined using the matchmaker function.

### Phylogeny of bat Cytochrome B gene

The evolutionary history of chiropteran cytochrome B was inferred by using the Maximum Likelihood method and JTT matrix-based model^83^. The tree with the highest log likelihood (−1568.10) is shown. The percentage of trees in which the associated taxa clustered together is shown next to the branches. Initial tree(s) for the heuristic search were obtained automatically by applying Neighbor-Join and BioNJ algorithms to a matrix of pairwise distances estimated using the JTT model, and then selecting the topology with superior log likelihood value. The tree is drawn to scale, with branch lengths measured in the number of substitutions per site. This analysis involved 5 amino acid sequences. Sequences of Cytochrome B were obtained from uniprot for each species: *Rhinolophus ferrumequinum* (accession O21298), *Eidolon helvum* (accession A0A7G3WDD6), *Rousettus leschenaultii* (accession B9VHX7), *Pteropus dasymallus* (accession Q9G6M3), and *Epomophorus gambianus* (accession G1C2G6). There were a total of 379 positions in the final dataset. Evolutionary analyses were conducted in MEGA X^82^.

### Production of Pseudotyped HNVpp-Rluc

HEK293T cells (4 × 10⁶) were seeded in collagen-coated 10-cm² dishes and transfected at 24hr post-seeding using BioT (Bioland Scientific) with 10 micrograms each of codon-optimized plasmids encoding homotypic HNV-F and RBP. At 16 HPT, cells were infected with VSV-ΔG-Rluc at a multiplicity of infection (MOI) of 10. Following two washes in PBS, fresh DMEM (10% FBS) supplemented with 1:10,000 anti-VSV-G monoclonal antibody (EB0010, Kerafast) was added. Supernatants were collected and clarified (1200 rpm, 5 min) at 48 HPI. GhVpp were concentrated by ultracentrifugation over a 20% sucrose cushion (pelleted using 25,000 rpm for 2hr at 4 °C) and resuspended in PBS. Infections were performed at a titer yielding at least a 3-log Rluc signal over Rluc background.

### HNVpp Neutralization Assay

Neutralization assays were conducted using titrated HNVpp-Rluc incubated with serial dilutions of polyclonal sera for 1 h at 37 °C. Polyclonal sera were heat-inactivated at 56 °C for 30 min prior to use. The mixtures were added to U-87 MG cells (3 × 10⁴ cells/well). At 24 h post-infection, cells were lysed (Promega, E2810), and Renilla luciferase activity was measured using a Cytation 3 (BioTek).

### Flow Cytometry-Based Surface Binding Assay

HEK293T cells (4 × 10⁶) were seeded in 6-well plates during 24h and transfected (BioT, Bioland Scientific) with 2 μg/well codon-optimized plasmids encoding HNV-F-AU1, or RBP-HA. At 24 h post-transfection, cells were harvested mechanically in PBS and optionally stained with Violet viability dye (1:10,000, ThermoFisher, L34955) for 20 min in the dark. Cells were washed in PBS, pelleted down and stained with primary antibodies diluted in FACS buffer (1x PBS, 15% FBS, 17 mM EDTA) for 1 h at 37 °C. Following two washes (FACS buffer), cells were incubated with Alexa-647-conjugated secondary antibodies (1:2000) for 1 h at room temperature in the dark. Washed twice, and resuspended in PBS. HNV glycoproteins were stained independently to measure expression using 1:1000 AU1 (Sigma Aldrich, F3165) or HA (Novusbio, NB600-363) mAb with corresponding secondary antibody Alexa 647. Flow cytometry was performed on an Attune NxT, using a no-primary control for gating. Data were analyzed in FlowJo v10.9.0, and GMFI values were plotted in GraphPad Prism v9.0.0. GMFI was normalized to expression measured with extracellular tag.

