## Supplemental Materials for "*De novo* recovery of Ghana virus, an African bat Henipavirus, reveals differential tropism and attenuated pathogenicity compared to Nipah virus"

### Supplemental Figures

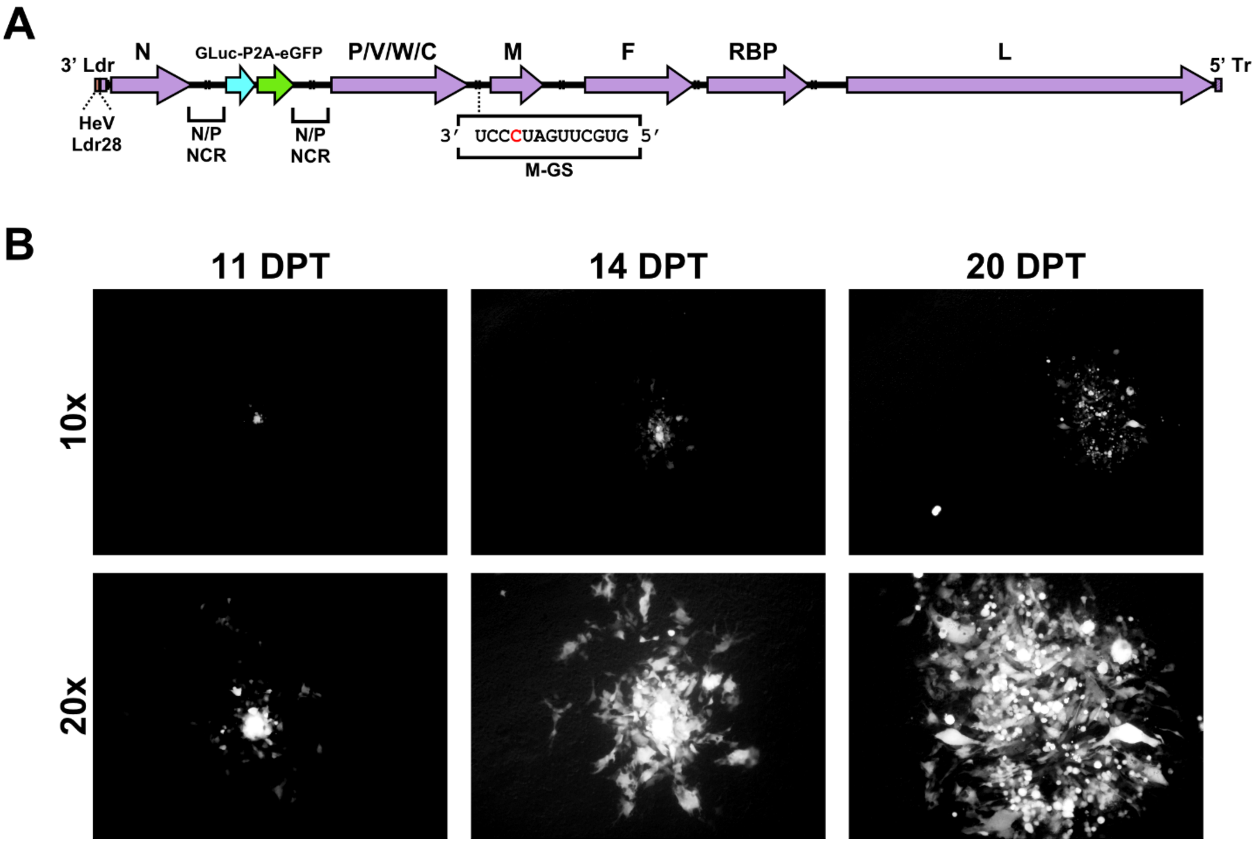


**Figure S1. Rescued rGhV M74a relies upon cell-to-cell spread in culture. (A)** Schematic of the rGhV genome. A *Gaussia* luciferase eGFP reporter gene cassette is accommodated by duplication of the NCR between the N and P genes. The unmapped 28 nucleotides of the GhV M74a genomic promoter sequence were replaced with the homologous genomic promoter sequence of Hendra virus (HeV Ldr28). The non-canonical M-GS of GhV strain M74a is shown in the inset box. **(B)** Microscopy depicting the spread of a GFP+ foci in the rGhV rescue plate over time. Foci were monitored between 11 and 20 days post transfection (DPT) for signs of spread. Images depict two magnifications of the same foci at each timepoint.

### Supplemental Figures

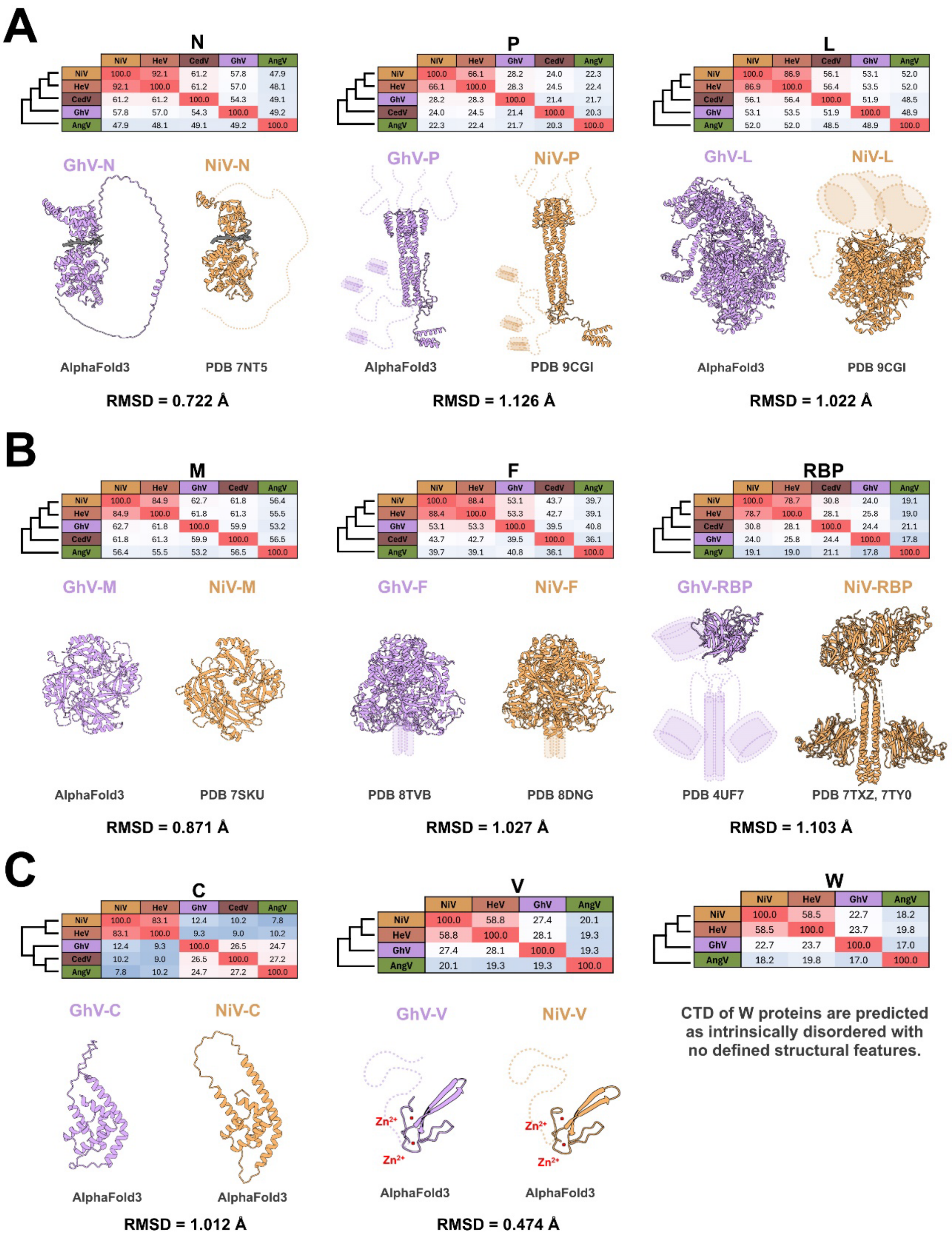


**Figure S2. *In silico* analysis of all proteins encoded by rGhV M74a.** In silico analyses of **(A)** the GhV replicase, consisting of GhV-N, GhV-P, and GhV-L, **(B)** the viral proteins required for particle formation and spread, GhV-M, GhV-F, and GhV-RBP, and **(C)** accessory proteins including GhV-C, GhV-V, and GhV-W. All encoded proteins in rGhV M74a were subject to amino acid sequence alignment with other bat-borne HNVs to generate phylogenies and percent identity matrices. Phylogenetic trees represent relatedness and are not to scale. Sequence

### Supplemental Figures

alignments were achieved using Clustal Omega, and phylogenies were constructed using MEGA X. Percent identity matrices are colored such that red is highly conserved and blue is less conserved. Structural comparisons were made in ChimeraX by comparing available structures for NiV proteins with available structures or AlphaFold3 models for GhV homologs. Root mean square deviation (RMSD) values were determined in ChimeraX using the matchmaker analysis.

### Supplemental Figures

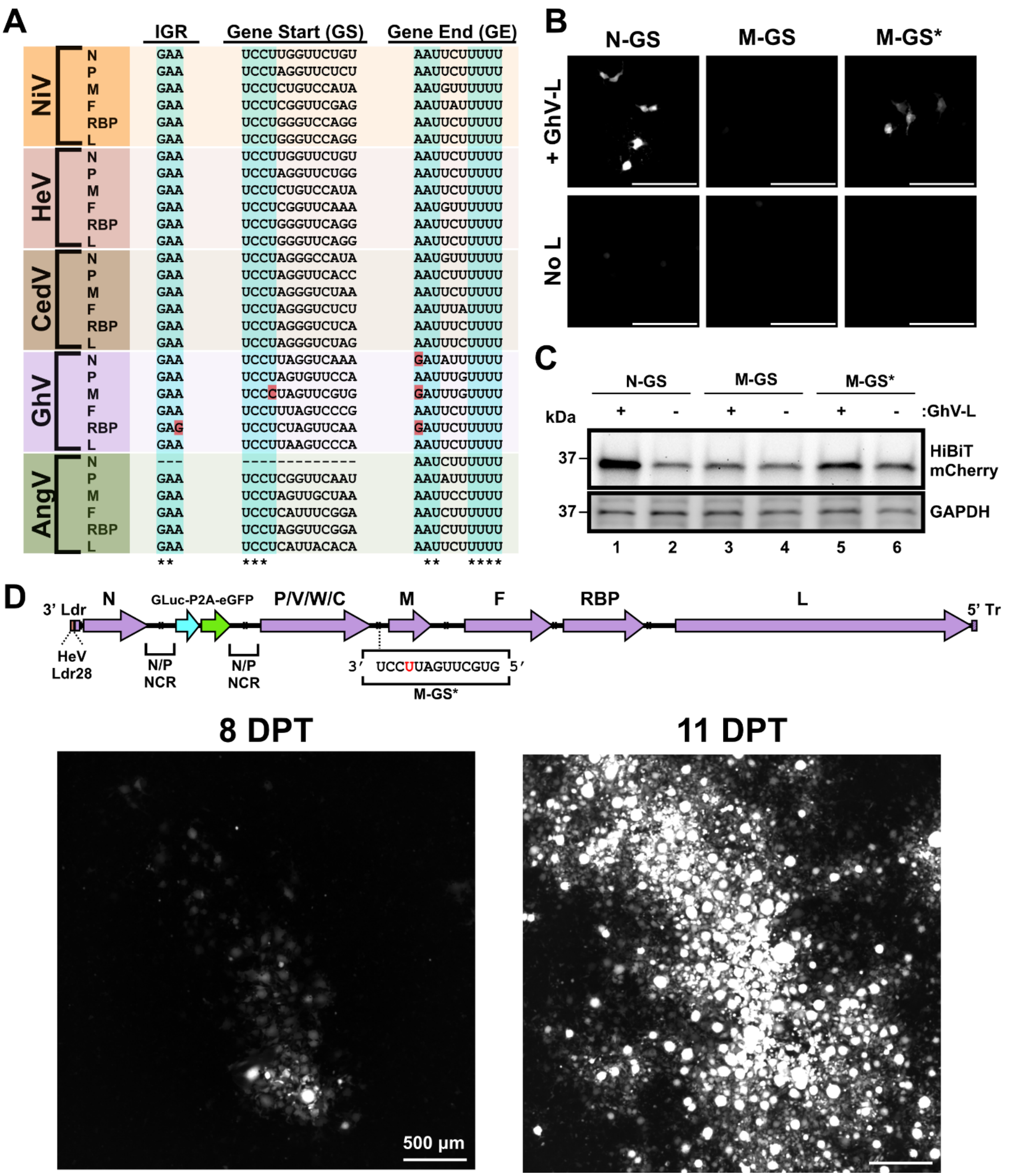


**Figure S3. The M-GS of GhV M74a is noncanonical, and does not drive detectable protein expression in GhV minigenome. (A)** Sequence alignment of all bat-borne HNV intergenic region (IGR), gene start (GS), and gene end (GE) sequences. Highly conserved motifs are highlighted in blue, and exceptions to these consensus motifs are colored in red. Absolutely conserved sequences are denoted with an asterisk (*) below the alignment. **(B)** Microscopy of mCherry positive events from each condition in **Figure 1C**. Scale bar represents 150 micrometers. **(C)** Western blot analysis of HiBiT-mCherry reporter gene expression from minigenome transfections in the presence or absence of GhV-L. Samples are matched with the experiment in **Figure 1C** and **S3B**. In this minigenome reporter construct, a 'background' HiBiT-mCherry band is often seen in the absence of L due to leaky expression of the T7pol-driven transcripts. In functional minigenomes, vRdRp-mediated increases in HiBiT-mCherry expression over background are observed in the presence of GhV-L. **(D)** Microscopy of a GFP+ foci in the rescue well of $rGhV_{M\text{-}GS*}$ imaged at 8 DPT and 11 DPT. Scale bar represents 500 micrometers.

### Supplemental Figures

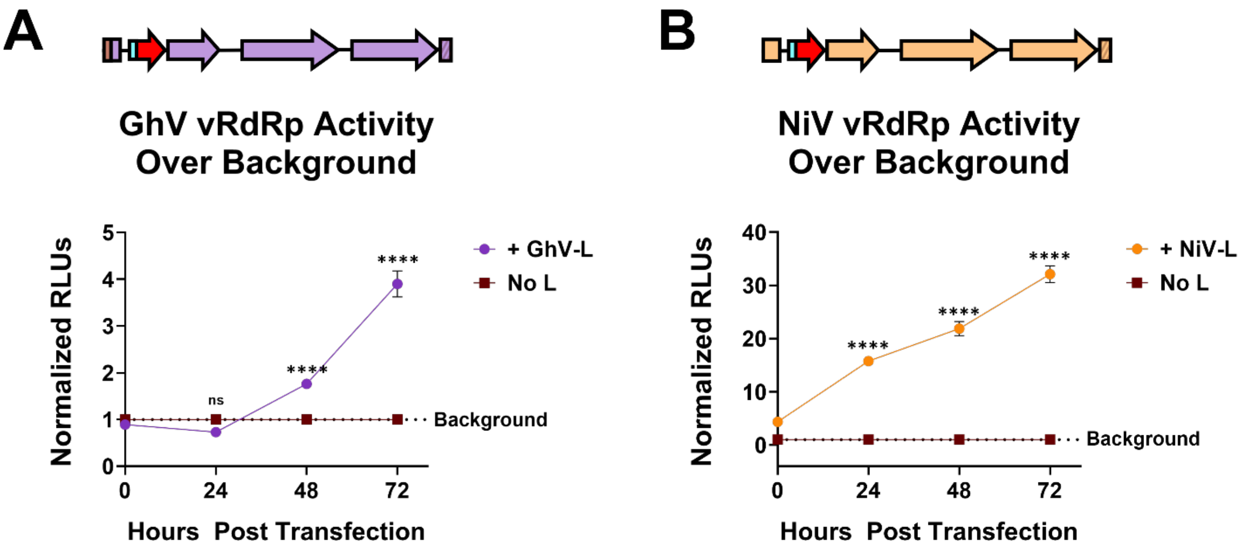


**Figure S4. Ghana virus and Nipah virus minigenome kinetics.** Signal over background in cells transfected with **(A)** the GhV TC-tr minigenome or **(B)** the NiV TC-tr minigenome in the presence or absence of respective L protein over time. RLUs from cells in the presence of HNV-L were normalized to respective RLUs generated by each minigenome in the absence of HNV-L at each timepoint. Statistical significance was assessed using a two-way ANOVA analysis in GraphPad Prism to compare counts or normalized RLUs in the presence of HNV-L with respective counts or normalized RLUs in the absence of HNV-L. All experiments were conducted in biological quadruplicate. Error bars depict standard deviation. For all graphs: ns, $P > 0.05$; *, $P \leq 0.05$; **, $P \leq 0.01$; ***, $P \leq 0.001$; ****, $P \leq 0.0001$.

**Supplemental Figures**

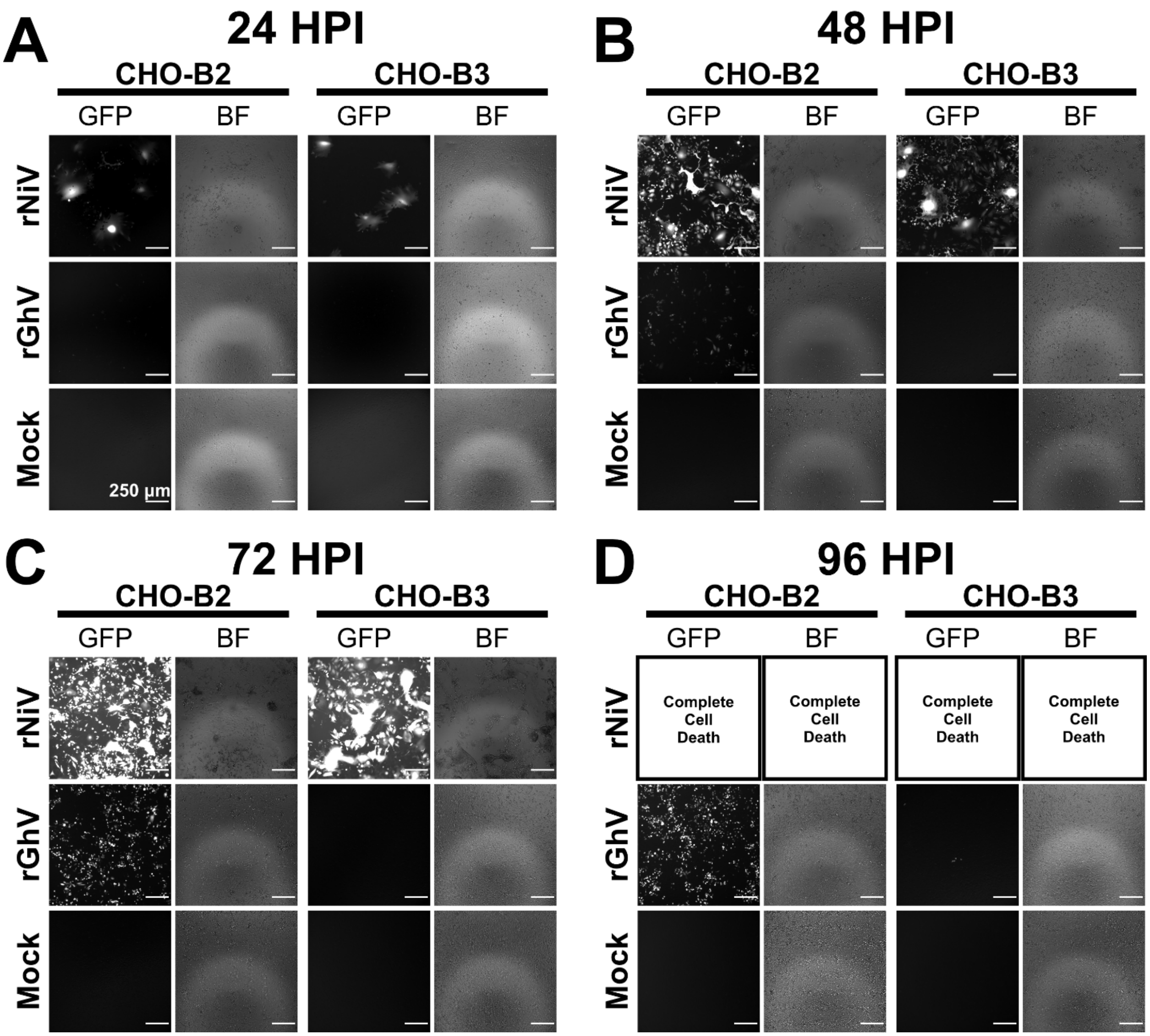


**Figure S5. Microscopy of rGhV$_{M\text{-}GS^*}$ and rNiV growth in CHO-B2 and CHO-B3 cells over time.** CHO-B2 and CHO-B3 cells were infected with either rNiV or rGhVM-GS* at an MOI of 0.05 and imaged at **(A)** 24 HPI, **(B)** 48 HPI, **(C)** 72 HPI, and **(D)** 96 HPI. Images denote GFP signal or brightfield (BF). Scale bars demonstrate 250 micrometers. Images are matched to the experiment in **Figure 1K**.

### Supplemental Figures

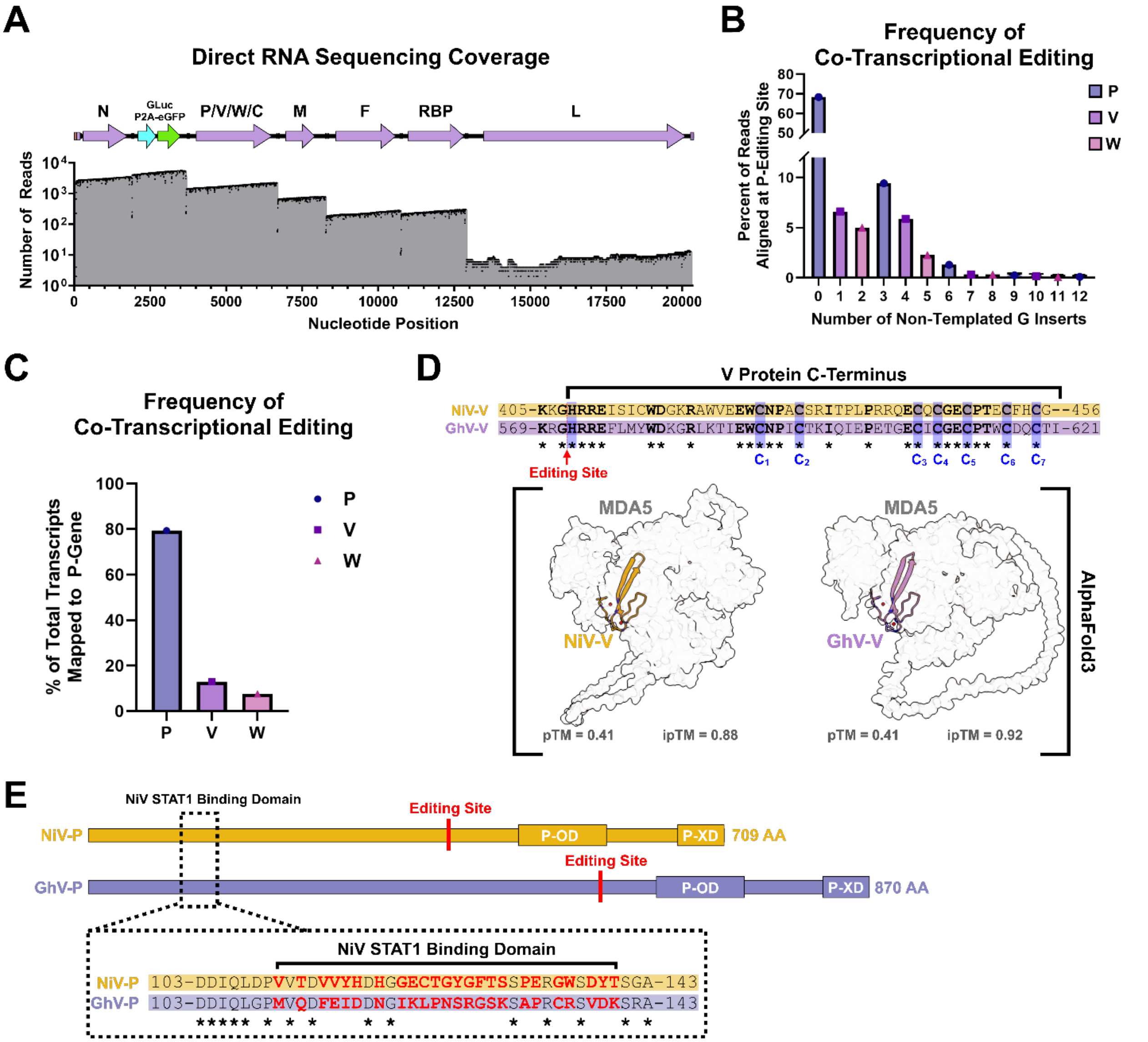


**Figure S6. Direct RNA sequencing of rGhV$_{M\text{-}GS*}$ infected CHO-B2 cells reveals co-transcriptional editing of the GhV-P gene transcripts. (A)** Coverage of long-read Nanopore direct RNA sequencing reads aligning to the rGhV$_{M\text{-}GS*}$ genome. **(B)** Frequency of co-transcriptional editing at the level of G insertions. P editing was measured using mPileup in Samtools, selecting reads with inserts of G only. **(C)** Data in (B) plotted as the respective products P, V, and W. **(D)** Sequence alignment of the C-terminus of NiV-V (orange) and GhV-V (purple) demonstrates conservation of the seven cysteines (highlighted in blue) required for zinc-binding. Below the alignment are AlphaFold3 predictions of the C-terminus of either NiV or GhV interacting with human MDA5; respective pTM and iPTM scores are listed below each model. **(E)** Depiction of the NiV-P and GhV-P proteins (to scale) and their domain organization. The described NiV STAT1-binding motif is highlighted, and sequence alignments show poor conservation at this site with mismatches in bold red. For all alignments, an * indicates a conserved residue between NiV and GhV.

### Supplemental Figures

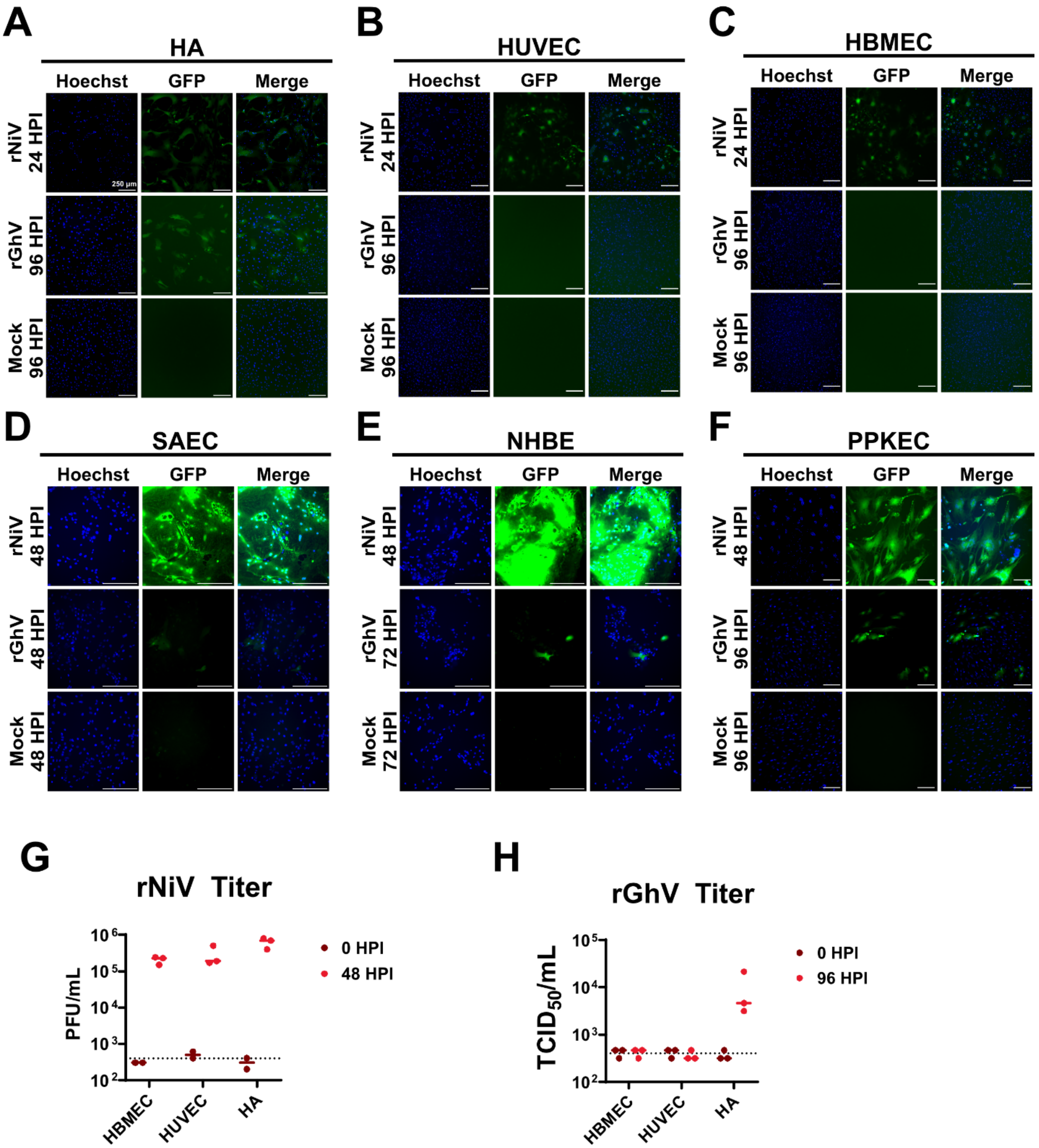


**Figure S7. Infection of primary human and porcine cells with rGhV$_{M\text{-}GS*}$ and rNiV.** Microscopy depicting rNiV and rGhV$_{M\text{-}GS*}$ infection of **(A)** HAs, **(B)** HUVECs, **(C)** HBMECs, **(D)** SAECs, **(E)** NHBEs, and **(F)** PPKECs. Timepoints of each image are denoted to the left of each image panel. Hoechst stain is colored in blue while GFP is colored in green. For all images, the scale bar is 250 micrometers. Virus in the supernatant of infected HBMEC, HUVEC, and HAs collected at 0 HPI and at the respective endpoint was titrated for **(G)** rNiV and **(H)** rGhV$_{M\text{-}GS*}$ to determine if infection was productive.

### Supplemental Figures

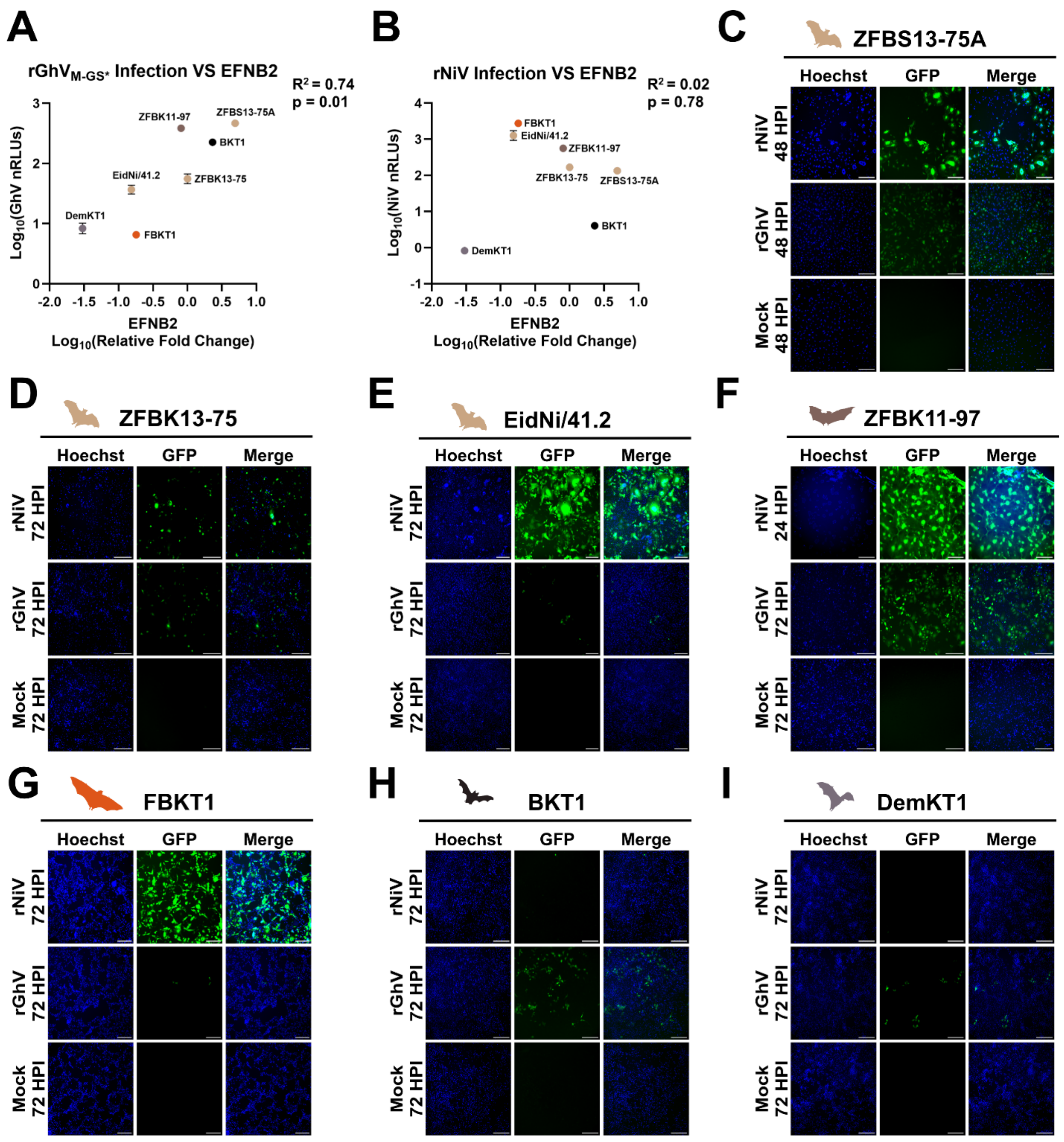


**J**

**EFNB2 G-H Loop**

| | |
|---|---|
| *H. sapiens* | FQEFSPNLWGLEFQKNKDYY |
| *P. alecto* | FQEFSPNLWGLEFQKNKDYY |
| *E. helvum* | FQEFSPNLWGLEFQKNKDYY |
| *R. leschenaultii* | FQEFSPNLWGLEFQKNKDYY |
| *R. ferrumequinum* | FQEFSPNLWGLEFQKNKDYY |
| | ******************** |

**K**

**EFNB3 G-H Loop**

| | |
|---|---|
| *H. sapiens* | FQEYSPNLWGHEFRSHHDYY |
| *P. alecto* | FQEYSPNLWGHEFRSHHDYY |
| *R. leschenaultii* | FQEYSPNLWGHEFRSHHDYY |
| *R. ferrumequinum* | FQEYSPNLWGHEFRSHHDYY |
| | ******************** |

**Figure S8. Infection of immortalized bat cell lines with rGhV$_{M-GS*}$ and rNiV.** Correlation plots comparing relative expression of EFNB2 and peak nRLUs produced in each bat cell line for **(A)** rGhV$_{M-GS*}$ and **(B)** rNiV. RT-qPCR was used to determine the relative expression of EFNB2. Expression of EFNB2

### Supplemental Figures

was first determined by respective comparison with GAPDH before then normalizing to the expression in ZFBK13-75. Pearson correlation coefficients ($R^2$) and p-values were calculated using GraphPad Prism. Correlations were computed between EFNB2 relative expression ($log_{10}$ fold change) and log-transformed nRLUs. Microscopy depicting rNiV and $rGhV_{M-GS*}$ infection of **(C)** ZFBS13-75A, **(D)** ZFBK13-75, **(E)** EidNi/41.2, **(F)** ZFBK11-97, **(G)** FBKT1, **(H)** BKT1, and **(I)** DemKT1 cells. All cells were infected at an MOI of 0.05 as titered on CHO-B2. Timepoints of each image are denoted to the left of each image panel. Hoechst stain is colored in blue while GFP is colored in green. For all images, the scale bar is 250 micrometers. Sequence alignment of the G-H loop of **(J)** EFNB2 or **(K)** EFNB3 from respective species. Alignments were conducted using ClustalOmega in MegaX software.

### Supplemental Figures

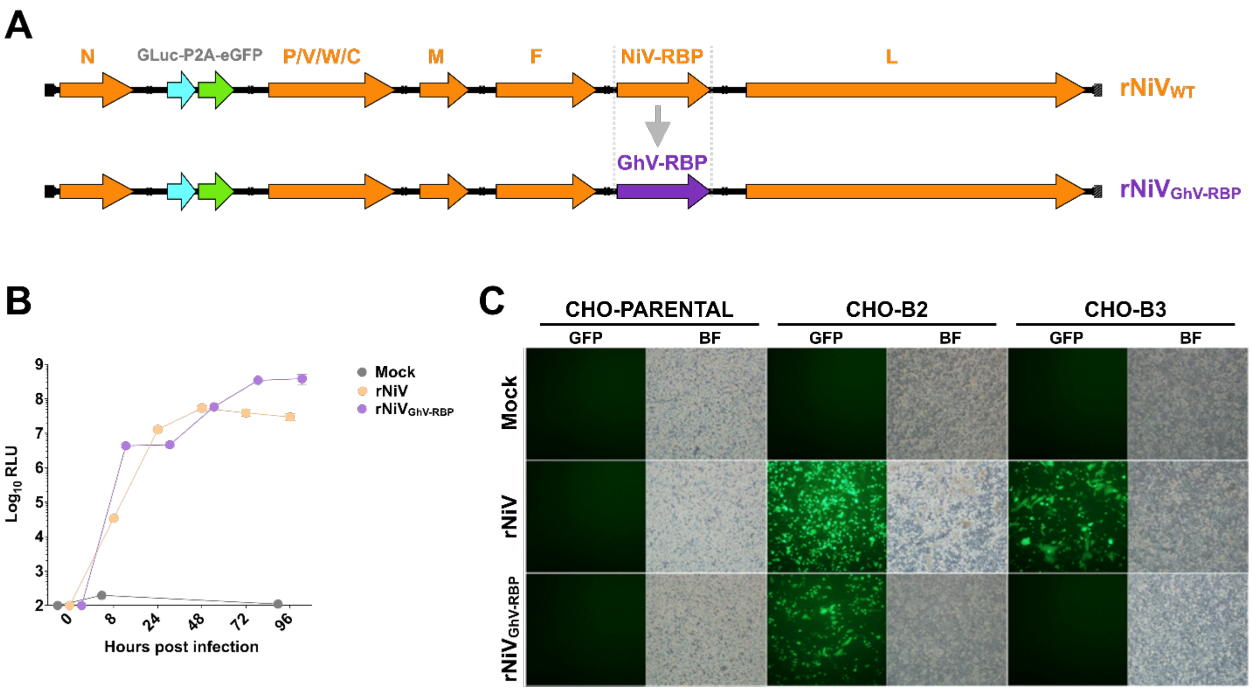


**Figure S9. Design and rescue of a rNiV chimera encoding the GhV-RBP. (A)** Schematic denoting the genome design of $rNiV_{GhV\text{-}RBP}$ relative to the parental, WT rNiV construct. **(B)** Growth kinetics of rNiV and $rNiV_{GhV\text{-}RBP}$ in Vero CCL81 cells, measured by *Gaussia* luciferase assay. **(C)** Comparative infection of CHO, CHO-B2, and CHO-B3 cells with either rNiV or $rNiV_{GhV\text{-}RBP}$.

### Supplemental Figures

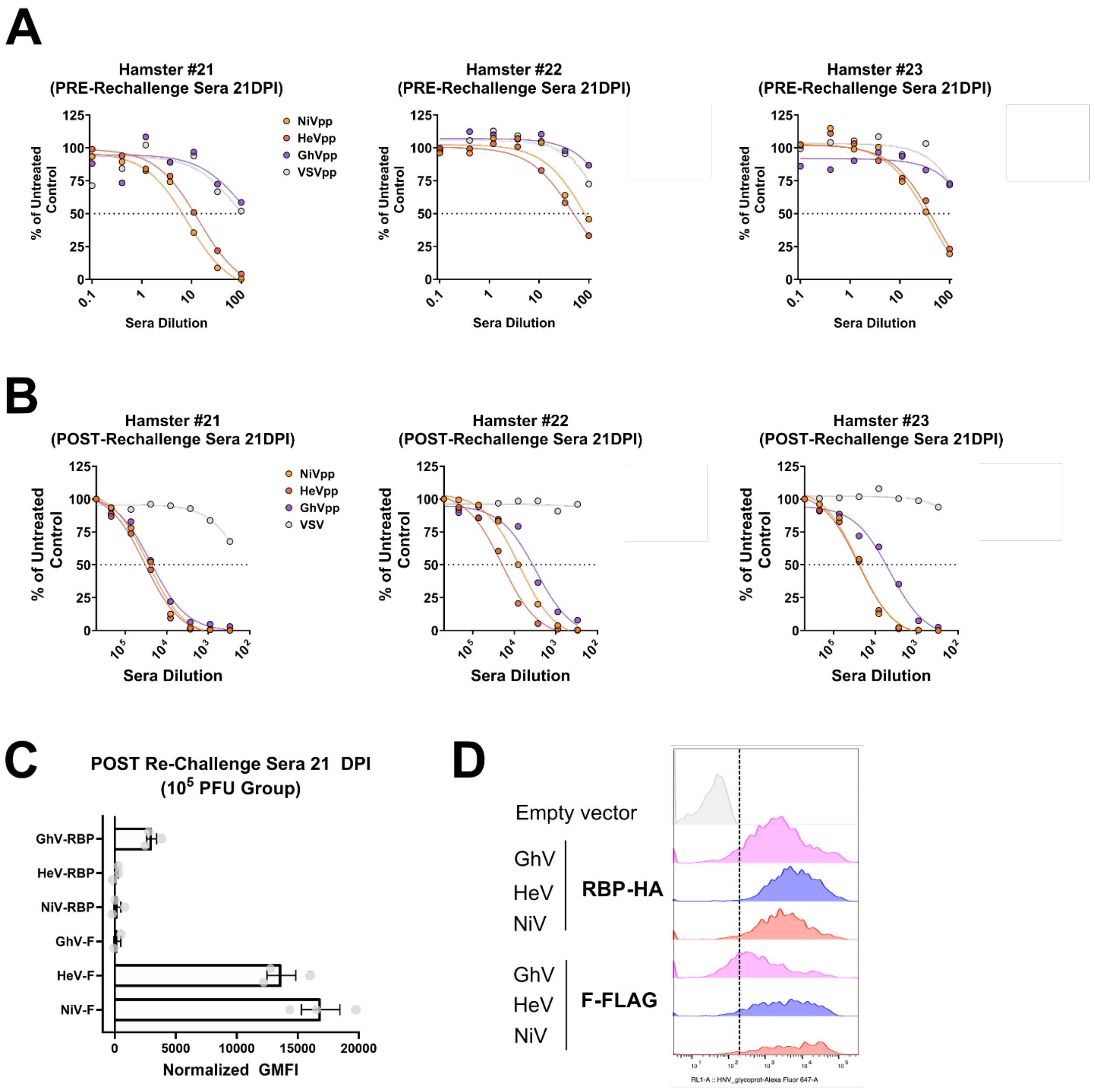


**Figure S10. Characterization of PRE and POST rechallenge sera from hamsters challenged initially with $10^5$ PFU of rNiV$_{GhV\text{-}RBP}$. (A)** Hamster sera collected at 21 DPI after the initial challenge with $10^5$ PFU of rNiV$_{GhV\text{-}RBP}$ pre-rechallenge was used in neutralization assay to measure inhibition of VSV pseudotyped with either NiV-F/RBP, HeV-F/RBP, GhV-F/RBP, or VSV-G as a control. **(B)** Hamster sera collected at 21 DPI after the rechallenge with $10^6$ PFU of WT rNiV was used in neutralization assay to measure inhibition of VSV pseudotyped with either NiV-F/RBP, HeV-F/RBP, GhV-F/RBP, or VSV-G as a control. **(C)** Flow cytometry was used to determine the ability of antibodies in post-rechallenge hamster sera to bind to HNV envelope proteins. For (A) and (B), RLUs were normalized to untreated control for each virus. For (C), GMFI was normalized to expression of each viral protein, shown in **(D)**.
